# Phosphorylation remodels the mitotic centrosome matrix to generate bipartite γ-tubulin complex docking sites

**DOI:** 10.1101/2025.11.20.689565

**Authors:** Midori Ohta, Orie Arakawa, Yajie Gu, Wanying Tian, Kevin D. Corbett, Arshad Desai, Karen Oegema

## Abstract

Mitotic centrosomes consist of centrioles surrounded by a proteinaceous matrix that docks and activates γ-tubulin complexes (γTuCs) to nucleate microtubules for spindle assembly. During mitotic entry, phosphorylation at centrosomes remodels CDK5RAP2 family matrix proteins to generate γTuC docking sites. We address the mechanism of this conversion using *C. elegans* SPD-5 as a model. We show that SPD-5 contains two regions, PRGB1 and PRGB2, that are each sufficient for Polo-Like Kinase 1 (PLK1) phosphorylation–regulated γTuC binding. We define key phosphosites in each region and uncover autoinhibition mediated by interactions within and between them. PRGB2 is dimeric and requires γTuCs containing the Mozart family microprotein MZT-1 for binding, whereas PRGB1 is monomeric and binds independently of MZT-1. Our results support PLK1 phosphorylation inducing a conformational change that enables MZT-1–dependent PRGB2 binding, which in turn relieves PRGB1 inhibition. Such a multi-step mechanism would ensure robust spindle assembly by restricting microtubule nucleation in space and time.

## Introduction

Centrosomes consist of a centriolar core surrounded by a pericentriolar material (PCM) matrix that nucleates and anchors microtubules (*1, 2*). During cell division in metazoans, centrosomes catalyze formation of the mitotic spindle (*3–9*). In preparation for mitosis, the PCM matrix increases in size and microtubule nucleation capacity in a process controlled by Polo-like Kinase 1 (PLK1) (*10–16*). Across systems, the PCM matrix is assembled primarily from CDK5RAP2 family proteins: CDK5RAP2 in humans, Centrosomin (Cnn) in *Drosophila*, and SPD-5 in *C. elegans* (*17*). In *Drosophila*, the PCM matrix expands via a phospho-regulated self-interaction between Cnn molecules in which the C-terminal region of one Cnn molecule interacts with an internal leucine zipper in a second Cnn molecule that has adjacent PLK1 phosphorylation sites that regulate assembly (*11, 18, 19*); a similar set of PLK1 sites controls assembly of *C. elegans* SPD-5 (*16*).

Complexes containing the specialized tubulin isoform γ-tubulin make a major contribution to microtubule nucleation at centrosomes (*20–22*). γ-tubulin is incorporated into small Y-shaped hetero-tetrameric complexes (γTuSCs) that contain two γ-tubulin molecules and two related scaffold proteins (GCP2 and GCP3 in humans). These γTuSCs can laterally associate to form larger soluble γ-tubulin ring complexes (γTuRCs) (*20, 22*). Work in *Drosophila* has suggested that γ-tubulin complexes are recruited to the centrosome by two main pathways (*23*). Preassembled γTuRCs can be brought to the PCM matrix by the centrosomal protein Spd-2 or the CDK5RAP2-like protein Cnn. The Spd-2 pathway can only recruit γTuRCs, whereas the Cnn pathway can also recruit smaller γTuSCs, which presumably laterally associate to form ring-shaped complexes on their docking sites (*20, 23–25*). Humans, *Xenopus*, and *Drosophila* have soluble γTuSCs and γTuRCs (*26–30*) and likely utilize both recruitment pathways. In contrast, *C. elegans* and budding yeast only have γTuSCs (*20, 31–33*) and depend solely on a Cnn-like mechanism for γ-tubulin recruitment.

The nucleating activity of γ-tubulin-containing complexes is thought to be activated upon recruitment to microtubule organizing centers such as the centrosome (*20, 22*). During mitosis in *C. elegans*, PLK1 phosphorylates SPD-5 on two sets of sites that independently regulate distinct functions: phosphorylation of the C-terminal half promotes expansion of the SPD-5-based PCM matrix while phosphorylation of the N-terminus converts it into a γ-tubulin complex docking site (*32*). In *Drosophila*, phosphorylation of the N-terminus of Cnn has similarly been proposed to generate γ-tubulin complex docking sites by relieving an autoinhibited state (*34*).

Early studies in fission yeast identified a conserved protein sequence shared by γ-tubulin complex docking proteins that was later named CM1 (Centrosomin Motif 1; (*35, 36*)). Work in human cells partitioned CM1 into two conserved regions, a stimulatory N-terminal region and an autoinhibitory C-terminal region. The N-terminal region of CM1 contains an ∼30aa peptide termed the γTuNA (γ-TuRC-mediated nucleation activator; (*37*)) that forms a small dimeric coiled-coil that can bind and activate γTuRCs (*37–41*). Structural work has shown that the γTuNA dimer forms a tripartite complex with the N-terminal helical domain (NHD) of GCP2 and a 3-helix Mozart family microprotein to form modules that promote interactions between adjacent γTuSCs in the γTuRC to bring them into a configuration that promotes microtubule nucleation (*41–43*). When N-terminal fragments of CDK5RAP2 are extended beyond the γTuNA to include the second conserved CM1 motif, which has been termed γTuNA inhibitor-In (γTuNA-In; (*44*)), they lose the ability to interact with γ-tubulin complexes. Collectively, these data suggest that γ-tubulin complex docking sites on PCM matrix molecules are held in an autoinhibited conformation that is relieved by phosphorylation, yet how the N-termini of CDK5RAP2-like proteins are restructured by phosphorylation to relieve autoinhibition and how they interact with γ-tubulin complexes to activate them remain open questions.

Here, we use the *C. elegans* PCM matrix protein SPD-5 as a model to determine how phosphorylation converts it into a γ-tubulin complex docking site. We find that this regulation is surprisingly complex. First, we show that the SPD-5 N-terminus contains two regions positioned on either side of an inhibitory region with homology to the γTuNA-In that are each sufficient for PLK1 phosphorylation-regulated γTuC binding *in vitro* and required for γTuC recruitment *in vivo*. These regions, which we name PRGB1 and PRGB2 (<u>P</u>hospho-<u>R</u>egulated <u>γ</u>TuC <u>B</u>inding) are mechanistically distinct in how they recognize γTuCs. In addition to positive phosphoregulation of each region, we uncover autoinhibition mediated by interactions within and between the two regions. We show that in the intact N-terminus, PLK1 phosphorylation induces a conformational change that enables MZT-1-dependent γTuC binding to PRGB2, which in turn relieves inhibition of PRGB1 by the γTuNA-In to permit its γTuC binding. Our findings reveal the multi-step conversion of a PCM matrix scaffold protein into a bipartite γTuC docking site, which we propose constrains microtubule nucleation to mitotic centrosomes to enable rapid and robust spindle assembly.

## Results

### SPD-5 fragments exhibit a complex pattern of phosphorylation-controlled binding to γ-tubulin complexes

During mitotic entry in the *C. elegans* embryo, the PCM matrix surrounding the centrioles, which is composed of the CDK5RAP2 family protein SPD-5, increases 5- to 10-fold in size in a process known as centrosome maturation (**Fig. 1A**; (*45–47*)). Centrosome maturation is driven by the mitotic kinase PLK1 (*10–16*) and the regulatory protein SPD-2 (homolog of human CEP192; (*48, 49*)). SPD-2, which has a mitotic docking site for PLK1, binds SPD-5 and delivers PLK1 to the PCM matrix (*50, 51*). PLK1 phosphorylates the central region of SPD-5 to promote SPD-5–SPD-5 interactions that drive matrix expansion and independently phosphorylates the SPD-5 N-terminus to transform it into a γ-tubulin complex (γTuC) docking site (**Fig. 1B**; (*16, 32*)). To understand how PLK1 phosphorylation converts the SPD-5 N-terminus into a γTuC docking site, we combined an *in vitro* reconstitution system for the interaction of γTuCs with the PLK1-phosphorylated SPD-5 N-terminus with *in vivo* analysis (*32*). In this reconstitution, the components of the *C. elegans* γTuC—γ-tubulin^TBG-1^, GCP2^GIP-2^, GCP3^GIP-1^, and the Mozart family protein MZT-1 (*31–33*)—are co-expressed in FreeStyle 293-F human cells and immunopurifed onto beads (**Fig. 1C**). The ability of SPD-5 fragments purified from bacteria to bind γTuC-coated beads in the presence or absence of PLK1 phosphorylation, is then assessed. In prior work, we showed that SPD-5 aa 1-473 exhibits robust PLK1 phosphorylation-dependent binding to γTuC-coated beads in this assay and identified two predicted PLK1 phosphosites (T178 and T198) important for binding (*32*). However, how PLK1 phosphorylation at these and other sites in the SPD-5 N-terminus stimulates γTuC binding has not been addressed.

**Fig. 1.**
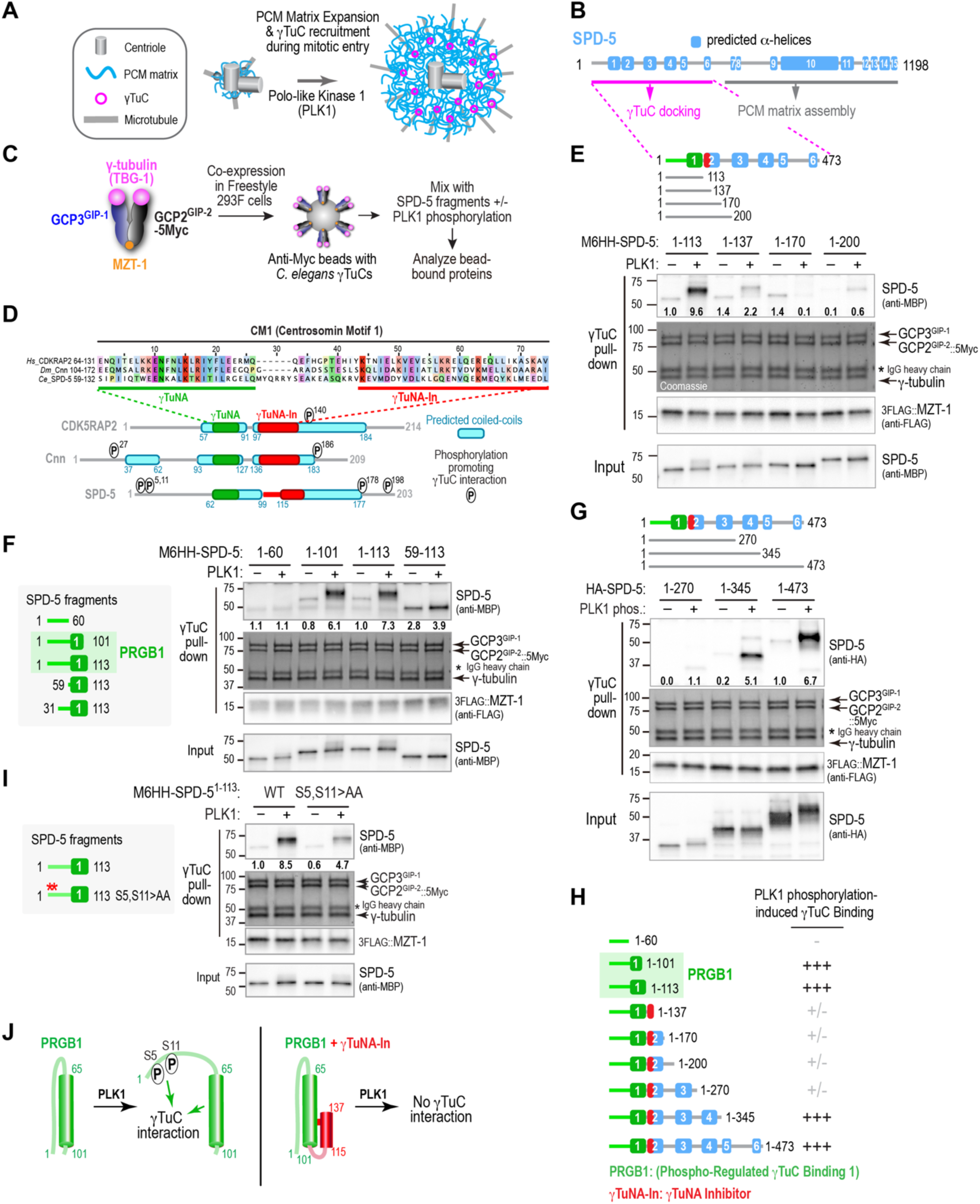
SPD-5 N-terminal fragments exhibit a complex pattern of PLK1 phosphorylation-dependent binding to γ-tubulin complexes. **(A)** Schematic illustrating how PLK1 kinase promotes centrosome maturation by phosphorylating the CDK5RAP2 family protein SPD-5. **(B)** SPD-5 schematic showing its fifteen predicted α-helices. Phosphorylation of its C-terminal half promotes matrix assembly, whereas phosphorylation converts its N-terminus into a docking site for γ-tubulin complexes (γTuCs). **(C)** Schematic of the assay to assess PLK1-dependent γTuC binding by SPD-5 fragments *in vitro*. γ-tubulin^TBG-1^, GCP2^GIP-2^-5Myc, GCP3^GIP-1^, and MZT-1 are co-expressed in Freestyle 293F cells, and complexes are immunoisolated onto Myc antibody-coated beads. SPD-5 fragments, purified from bacteria and preincubated with or without PLK1 in the presence of ATP, are mixed with γTuC-immobilized beads and bead-bound proteins are analyzed. **(D)** ClustalWS alignment of the CM1 region from three CDKRAP2 family proteins, human CDK5RAP2, *Drosophila* Cnn and *C. elegans* SPD-5. Schematics of the N-termini of the three proteins highlight predicted coiled-coils (PCOILS 21 aa window rounded score ≥0.5), the location of the sequences aligned with the γTuNA (***green***) and γTuNA-In (***red***) and previously identified sites whose phosphorylation promotes γTuC binding ((*32, 34, 65*); SPD-serine 5 and 11 identified in this study. **(E-G & I)** Binding assays, conducted as outlined in (*C*), with γTuC-coated beads and MBP-6His-HA (M6HH)-tagged (*E,F,I*) or HA-tagged (*G*) SPD-5 fragments, preincubated with or without PLK1 as indicated. SPD-5 and MZT-1 were analyzed by immunoblotting using the indicated antibodies; γ-tubulin, GCP2^GIP-2^, and GCP3^GIP-1^ were detected by Coomassie staining. Numbers below the SPD-5 fragment bands indicate band intensity relative to the SPD-5 1-113 (*E,F,I*) or 1-473 (*G*) in the absence of PLK1 phosphorylation. Asterisk indicates the IgG heavy chain of the anti-Myc antibody used for the IP. **(H)** Schematic of the SPD-5 fragments used in (*E-G*) along with a summary of their PLK1 phosphorylation induced γTuC-binding. **(J)** Speculative model for how PLK1 phosphorylation impacts γTuC binding by fragments containing PRGB1 (*left*) or PRGB1 plus the γTuNA-In (*right*).

Prior work on γ-tubulin complex docking sites in CDK5RAP2 family proteins has focused on their CM1 regions (*22, 35–43*). Sequence alignments of SPD-5’s CM1-like region with the CM1 regions of *Drosophila* Cnn and human CDK5RAP2 (**Fig. 1D**) suggest that the *C. elegans* sequence has two adjacent regions whose sequence can be aligned with the previously described γTuNA (aa 59-84; (*37*)) and γTuNA inhibitor (γTuNA-In; aa 102-132; (*44*)) motifs. However, the putative SPD-5 γTuNA exhibits sequence divergence in key positions conserved in other family members, particularly the lysine and phenylalanine in the KENF motif (**Fig. 1D**), which motivated us to functionally test whether this region mediates binding to the γTuC in the *in vitro* reconstitution assay. Consistent with the alignment, a short N-terminal fragment of SPD-5 (aa 1-113) containing the predicted γTuNA, but not the γTuNA-In, bound to γTuC when phosphorylated by PLK1 (**Fig. 1E,F**); a slightly shorter fragment SPD-5 (aa 1-101) exhibited equivalent robust PLK1 phosphorylation-dependent binding (**Fig. 1F**), whereas a fragment lacking the predicted γTuNA did not (aa 1-60; **Fig. 1F**). Since the aa 1-101 or aa 1-113 fragments both exhibit phospho-regulated γTuC binding, we refer to them interchangeably as <u>P</u>hospho-<u>R</u>egulated <u>γ</u>TuC <u>B</u>inding <u>1</u> (PRGB1; **Fig. 1F**). In further agreement with the alignment, extending the N-terminal SPD-5 fragment beyond PRGB1 to include the region with homology to γTuNA-In (aa 1-137 and aa 1-170) strongly suppressed PLK1-phosphorylation-stimulated γTuC binding (**Fig. 1D,E**). Two additional N-terminal SPD-5 fragments (1–200, 1–270) that include the PLK1 phosphorylation sites previously shown to be important for γTuC binding *in vitro* and recruitment to centrosomes *in vivo* (T178 and T198; (*32*)) also failed to exhibit significant PLK1-phosphorylation-dependent binding (**Fig. 1E,G**). However, consistent with PLK1 phosphorylation-dependent binding being robust for the N-terminal half of SPD-5 (aa 1-473; (*32*)), further extension of the N-terminal region to aa 345 or aa 473 restored PLK1 phosphorylation-dependent γTuC binding (**Fig. 1G,H**).

The pattern of presence, loss, and then recovery of γTuC binding across the set of N-terminal SPD-5 fragments (**Fig. 1H**), suggests that the PLK1 phosphorylation-controlled mechanisms that convert the SPD-5 N-terminus into a γTuC docking site are complex. This pattern motivated us to undertake a detailed investigation of the conversion mechanism, starting with analysis of γTuC binding by PRGB1.

### Phosphorylation converts the region preceding the γTuNA from inhibitory to activating for γTuC binding

The results above indicated that PRGB1, which consists of the predicted γTuNA preceded by 60 N-terminal amino acids, exhibits robust PLK1 phosphorylation-dependent γTuC binding (**Fig. 1F,H**). Relative to PRGB1, fragments of PRGB1 that lacked the first 30 or 60 amino acids exhibited elevated binding when unphosphorylated and this binding was only slightly enhanced by PLK1 phosphorylation (SPD-5 aa 59-113, **Fig. 1F**; aa 31-113, **Fig. S1A**). This result has two implications. First, it suggests that the extreme N-terminus of SPD-5 (aa 1-60) inhibits phosphorylation-independent γTuC binding by the γTuNA and that this inhibition is released by PLK1 phosphorylation. Our prior work identified two candidate PLK1 target sites in the first 30 amino acids of SPD-5 that are broadly conserved across metazoans (Ser5 and Ser11; **Fig. S1B**; (*32*)). Mutating Ser5 and Ser11 to alanines reduced the ability of PLK1 to stimulate γTuC binding by PRGB1 (**Fig. 1I**), suggesting that phosphorylation of these sites helps to relieve inhibition of the γTuNA by the aa 1-60 region (**Fig. 1J**). Second, the fact that PLK1-phosphorylated PRGB1 binds significantly better to γTuCs than the smaller fragment containing only the γTuNA (**Fig. 1F**), suggests that phosphorylation of aa 1-60 contributes positively to γTuC binding by PRGB1 (**Fig. 1J**). Fragments that contain PRGB1 followed by the γTuNA-In (SPD-5 aa 1-137 or 1-170) cannot be activated by phosphorylation (**Fig. 1E**), indicating that addition of this predicted α-helical region inhibits γTuC binding by the γTuNA in a way that renders PRGB1 unable to be activated by PLK1 phosphorylation (**Fig. 1J**).

To determine whether the *in vitro* results align with the *in vivo* requirements for γTuC recruitment to centrosomes, we imaged embryos expressing γ-tubulin::mCherry and RNA-resistant single-copy transgene-encoded WT or mutant versions of GFP::SPD-5, following endogenous SPD-5 depletion (**Fig. 2A-C**). The ability of the SPD-5 matrix to assemble around centrioles during mitotic entry was not altered by deletion of the core γTuNA homology region (Δ61-93), truncation of the first 60aa (Δ60), or mutation of Ser5 and Ser11 to alanines (**Fig. 2A,B**). In contrast, all three of these mutations led to a similarly strong reduction in γTuC recruitment (**Fig. 2A,B**). These results confirmed the importance of the γTuNA homology region and of the first 60aa of SPD-5 in γTuC recruitment *in vivo*. They also support the conclusion from the biochemical analysis that phosphorylation of the 1-60 region, which includes Ser5 and Ser11, not only alleviates γTuNA inhibition but also contributes positive affinity to γTuC binding (**Fig. 1F,J**).

**Fig. 2.**
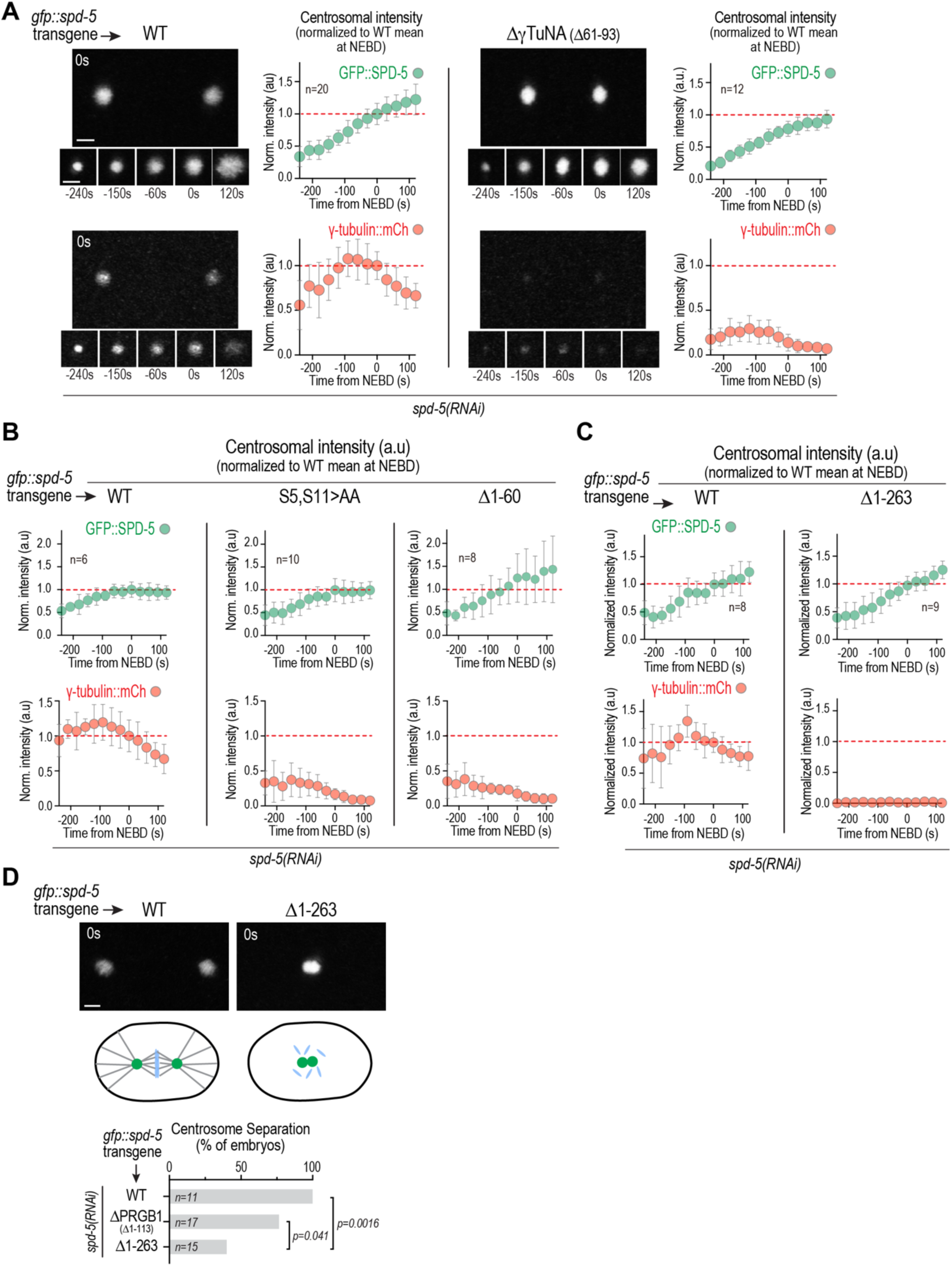
Impact of deletions in the SPD-5 N-terminus on γTuC recruitment to centrosomes. **(A-C)** Images and quantification (*A*) or quantification only (*B,C*) of centrosomal fluorescence intensity over time for γ-tubulin::mCherry and the indicated GFP::SPD-5 variants after depletion of endogenous SPD-5 by RNAi. Centrosomal fluorescence was normalized by dividing by the mean at nuclear envelope breakdown (NEBD) for WT GFP::SPD-5. The means for the WT control at NEBD are marked with a red dashed line. Error bars are the standard deviation (SD). *n* is the number of centrosomes imaged for each condition. **(D)** (*top*) Images of an embryo expressing WT GFP::SPD-5 with separated centrosomes and one expressing GFP::SPD-5 Δ1-263 that failed to separate its centrosomes. (*middle*) Schematics illustrating the location of the centrosomes and status of the spindle in the imaged embryos. (*bottom*) Graph plots the percentage of embryos with separated centrosomes expressing each of the indicated GFP::SPD-5 variants. Endogenous SPD-5 was depleted by RNAi in all conditions; *n* is the number of imaged embryos. Scale bars are 2 µm. P-values are one sided Fisher’s Exact Test. Note that separate WT controls are shown for (*A*), (*B*) and (*C*) because the data were collected at different times on different microscopes.

### Identification of a second region in the SPD-5 N-terminus that is sufficient for PLK1-regulated γTuC binding

Given the emphasis on the γTuNA as the primary recruiter of γTuCs, we anticipated that γTuC recruitment would be abolished when the γTuNA was absent in SPD-5. However, residual γ-tubulin was still detected at centrosomes in embryos expressing GFP::SPD-5 lacking the γTuNA (**Fig. 2A**). In contrast, in embryos expressing a larger N-terminal truncation of SPD-5 (Δ1-263), which also assembled normally around centrioles, γ-tubulin fell below the level of detection (**Fig. 2C**). We additionally observed that while the two centrosomes succeeded in separating from each other in the majority of cases in embryos expressing SPD-5 lacking PRGB1 (**Fig. 2D**; Δ1-113), suggesting that they retained partial ability to nucleate microtubules, centrosomes failed to separate in the majority of embryos with the larger aa 1-263 deletion (**Fig. 2D**; Δ1-263). These observations hinted that the SPD-5 N-terminus may contain a second, PRGB1-independent, means of interacting with γTuCs. Immediately after PRGB1 is α-helix 2 (aa 118-166; **Fig. 1D**). Addition of the putative γTuNA-In (SPD-5 aa 1-137), or all of predicted helix 2 (SPD-5 aa 1-170), to PRGB1 strongly suppressed PLK1 phosphorylation-stimulated binding to γTuCs (**Fig. 1E**), whereas phosphorylation-stimulated binding was restored by further extension of the N-terminus to aa 345 (**Fig. 1G**). This suggests that, in addition to a potential PRGB1-independent γTuC-recruitment module, elements of the SPD-5 N-terminus distal to PRBG1 and the γTuNA-In are also required to alleviate the potent inhibition of PRGB1 by the γTuNA-In that otherwise renders it impervious to PLK1 phosphoregulation.

To understand the roles of the region of SPD-5 distal to PRGB1 and the γTuNA-In, we analyzed γTuC binding by a series of N-terminal truncations of SPD-5 aa 1-473. SPD-5 fragments lacking PRGB1 (aa 101-473) or PRGB1 and the γTuNA-In (aa 135-473) exhibited robust PLK1 phosphorylation-dependent γTuC binding (**Fig. 3A**). In contrast, a shorter fragment (aa181-473) that lacked all of the predicted α-helix 2 failed to bind γTuCs in the presence or absence of PLK1 (**Fig. 3A**). These results suggest that aa135-180 is an essential part of a second, PRGB1-independent interface with γTuCs in SPD-5 (**Fig. 3A**). Consistent with the idea that the distal region of the SPD-5 N-terminus has dual roles in binding γTuCs and also in disengaging PRGB1 from the γTuNA-In to allow it to bind γTuCs, GFP-tagged Δ135-180 SPD-5, while assembling normally around centrioles, reduced γTuC recruitment to near background levels *in vivo* (**Fig. 3B**). Biochemical analysis revealed that SPD-5 aa135-345 exhibited robust PLK1 phosphorylation-dependent γTuC binding, whereas smaller fragments lacking either the N-terminal or C-terminal portions of the aa 135-345 region did not (**Fig. 3C**). Thus, both aa 135-180, which contain the portion of α-helix 2 after the γTuNA-In, and aa 200-345, which contain predicted α-helices 3 and 4, are required for phosphorylation-dependent γTuC binding. We note that aa 200-345 is also critical to alleviate inhibition of PRGB1 by the γTuNA-In (**Fig. 1E,G**). Collectively, these results define a second region of SPD-5 (aa135-345) that is capable of PLK1 phosphorylation-dependent γTuC binding on its own and is also required to alleviate inhibition of PRGB1 by the γTuNA-In. We named this region, which includes the portion of α-helix 2 after the <u>γ</u>TuNA-In along with predicted helices 3 and 4 PRGB2, (<u>P</u>hospho-<u>R</u>egulated <u>γ</u>TuC <u>B</u>inding <u>2</u>; **Fig. 3D**).

**Fig. 3.**
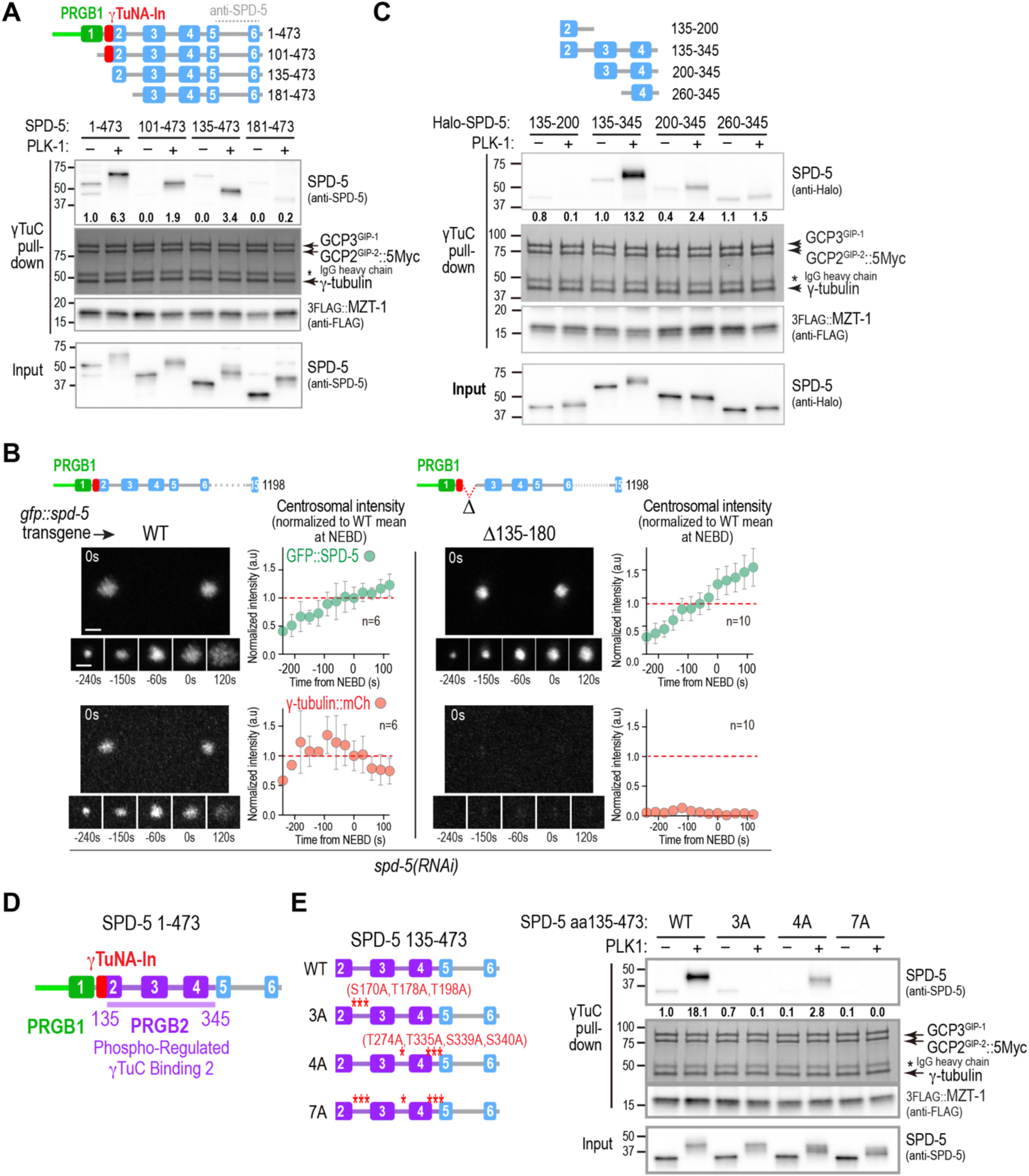
Identification of a second region in the SPD-5 N-terminus that is sufficient for PLK1-regulated γTuC binding. **(A,C)** Binding assays with γTuC-coated beads and the indicated untagged (*A*) or Halo-tagged (*C*) SPD-5 fragments, preincubated with or without PLK1 as indicated. SPD-5 and MZT-1 were analyzed by immunoblotting using the indicated antibodies; γ-tubulin, GCP2^GIP-2^, and GCP3^GIP-1^ were detected by Coomassie staining. Numbers below the SPD-5 fragment bands indicate band intensity relative to the SPD-5 1-473 (*A*) or SPD-5 135-345 (*C*) in the absence of PLK1 phosphorylation. Asterisk indicates the IgG heavy chain of the anti-Myc antibody used for the Myc IP. **(B)** Images and quantification of centrosomal fluorescence intensity over time for γ-tubulin::mCherry and the indicated GFP::SPD-5 variants after depletion of endogenous SPD-5 by RNAi. Centrosomal fluorescence was normalized by dividing by the mean at nuclear envelope breakdown (NEBD) for WT GFP::SPD-5. The means for the WT control at NEBD are marked with a red dashed line. Error bars are the standard deviation (SD). *n* is the number of centrosomes imaged for each condition. Scale bars are 2 µm. **(D)** Schematic showing the relative locations in the SPD-5 N-terminus of the two independent phospho-regulated γTuC binding elements, PRGB1 and PRGB2, and the intervening inhibitory γTuNA-In element. **(E)** Binding assays with γTuC-coated beads performed as in *(A,C)* for the indicated SPD-5 variants. Numbers below the SPD-5 fragment bands indicate band intensity relative to SPD-5 135-473 in the absence of PLK1 phosphorylation.

PRGB2 contains 3 PLK1 phosphorylation sites between helices 2 & 3 (S170, T178, T198), two of which (T178 and T198) were previously shown to be important for γTuC binding *in vitro* and *in vivo* (*32*). This region also contains 4 predicted PLK1 phosphorylation sites in the loops between predicted helices 3 & 4 and 4 & 5 (**Fig. 3E**; **Fig. S1B**). To analyze the significance of these phosphosites to PRGB2-mediated γTuC binding, we compared wildtype and alanine-mutated N-terminal fragments that lack PRGB1 (**Fig. 3E**). This analysis showed that residues S170, T178 and T198 are important for γTuC binding by PRGB2 and that the 4 other C-terminal PLK1 target sites also contribute to binding (**Fig. 3E**). We note that, although reduced in magnitude, a PLK1 phosphorylation-dependent shift in gel mobility was still detected for the 7A mutant, indicating the presence of additional PLK1 target sites beyond the ones mutated. Overall, these results identify PRGB2 as a second region of SPD-5 that exhibits PLK1 phosphorylation-dependent γTuC binding.

### PRBG2, but not PRGB1, requires the MZT-1 module for binding to γ-tubulin complexes

We next focused on assessing the elements of the γTuC that are important for binding to PLK1-phosphorylated PRGB1 and PRGB2. The *C. elegans* γTuC contains γ-tubulin, GCP2^GIP-2^, GCP3^GIP-1^, and MZT-1, the *C. elegans* homolog of Mozart1 (MZT1) (*20, 31–33*). MZT1 is a small, widely conserved microprotein composed of three α-helices that is required for γTuCs to interact with tethering proteins at MTOCs in plants, yeasts, and animal cells (*22, 52*). Structural work has shown that MZT1 intercalates with the N-terminal helical domain (NHDs) of the γTuSC subunit GCP3 and also with the γTuRC-specific components GCP5 and GCP6 (*22, 43, 53–59*). Vertebrates have a second Mozart protein, MZT2 that intercalates with the NHD of GCP2; in humans, the CDK5RAP2 γTuNA forms a parallel coiled-coil that interacts with the MZT2–GCP2 NHD module (*41–43*). In *C. elegans*, the single Mozart family protein, MZT-1, is required for γTuCs to dock onto the PLK1-phosphorylated SPD-5 matrix at centrosomes and for the SPD-5 N-terminus (aa 1-473) to bind γTuCs following PLK1-phosphorylation (*32, 33*). We thus focused on testing if the MZT-1-containing module is required for phosphorylated PRGB1 and PRBG2 to interact with γTuCs.

An AlphaFold 3 structural model of the *C. elegans* γTuC revealed a typical heterotetrameric small γ-tubulin complex (**Fig. 4A**). GCP3^GIP-1^, but not GCP2^GIP-2^, contains an N-terminal helical domain (NHD) that is predicted to form an intercalated structural module with MZT-1, which we refer to as the MZT-1 module (**Fig. 4A**). Consistent with this, expression of GCP3^GIP-1^ lacking its NHD in the reconstitution system resulted in γTuCs that did not incorporate MZT-1 (**Fig. 4B**). The MZT-1 module is connected to the rest of GCP3^GIP-1^ by an unstructured region and is not predicted to be specifically positioned relative to the rest of the γTuC (**Fig. 4A, Fig. S2A**); in addition, modeling 2 or more γTuCs did not yield high-confidence predictions that positioned the MZT-1 module relative to other complex subunits. Analyzing the ability of γTuCs that either contained or lacked the MZT-1 module to bind to PRGB1 and PRGB2 revealed that phosphorylated PRGB2 requires the MZT-1 module for γTuC binding, whereas PRGB1 does not (**Fig. 4C**, **Fig. S2B**). These results indicate that phosphorylated PRGB1 and PRGB2 engage γTuCs through distinct interfaces and can bind independently of one another when tested in isolation.

**Fig. 4.**
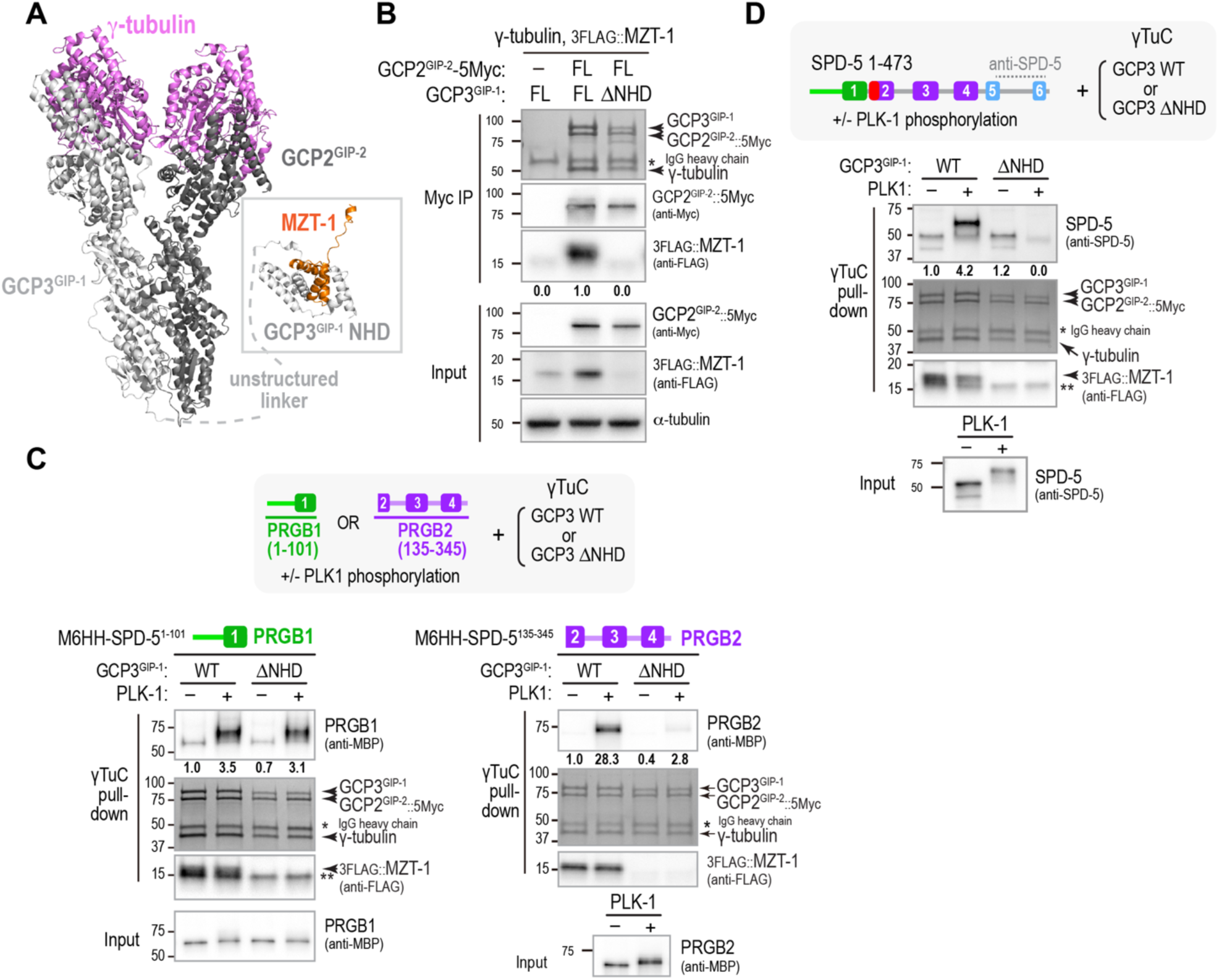
PRBG2, but not PRGB1, requires the MZT-1 module for binding to γ-tubulin complexes. **(A)** Alphafold 3 model of the *C. elegans* γTuC. The MZT-1 module comprised of MZT-1 and the GCP3^GIP-1^ N-terminal homology domain (NHD) is separated from the core tetrameric complex by an unstructured linker and is not positioned relative to the core complex. See also *Fig. S2A*. **(B)** Analysis of the composition of the γTuC when either full-length (FL) GCP3^GIP-1^ or GCP3^GIP-1^ with its N-terminal homology domain deleted (ΔNHD) was expressed. A condition in which GCP2^GIP-2^ with the Myc tag used for the immunoisolation omitted is shown as a control. MZT-1 and GCP2^GIP-2^ were detected by western blotting; γ-tubulin, GCP2^GIP-2^, and GCP3^GIP-1^ were detected by Coomassie staining. Numbers below the MZT-1 bands indicate band intensity relative to MZT-1 levels when FL GCP3^GIP1^ was expressed. Asterisk indicates the IgG heavy chain of the anti-Myc antibody used for the Myc IP. Note that MZT-1 is unstable in the absence of the GCP3^GIP-1^ NHD, so its levels are reduced in the input as well as in the immunoprecipitated complex. **(C,D)** Pull-down experiments with beads coated with γTuCs assembled with WT or ΔNHD GCP3^GIP-1^. Beads were incubated with the indicated SPD-5 fragments after preincubation with or without PLK1 as indicated. SPD-5 and MZT-1 were analyzed by immunoblotting using the indicated antibodies; γ-tubulin, GCP2^GIP-2^, and GCP3^GIP-1^ were detected by Coomassie staining. Numbers below the SPD-5 fragment bands indicate band intensity relative to PRGB1 (*C*, *left*), PRGB2 (*C*, *right*) or SPD-5 1-473 (*D*) in the absence of PLK1 phosphorylation. The single asterisk indicates the IgG heavy chain of the anti-Myc antibody used for the Myc IP. The double asterisk marks the location of a non-specific band.

In addition to its ability to bind γTuCs in a MZT-1 module-dependent manner, PRGB2 is required to alleviate the inhibition of PRGB1 by the γTuNA-In that otherwise renders it impervious to PLK1-dependent activation. Consistent with this, the ability of entire N-terminal region (aa 1-473) to bind γTuCs depends on the MZT-1 module that engages PRGB2 (**Fig. 4D**). These data indicate that when PRGB2 is unable to interface with the MZT-1 module, PRGB1 remains unable to form its independent interface with the γTuC, potentially due to an inability to disengage from the γTuNA-In.

### PRGB2 dimerizes the SPD-5 N-terminus and undergoes a conformational change upon PLK1 phosphorylation

We next focused on addressing how phosphorylation of PRGB2 promotes γTuC interaction. PRBG2 is comprised of predicted coiled-coils, raising the possibility that coiled-coil-mediated dimerization may be important for PRGB2 γTuC binding. To address this possibility, we analyzed purified SPD-5 fragments using size exclusion chromatography coupled with multi-angle light scattering (SEC-MALS) (**Fig. 5A**, **Fig. S3A,B**). The largest SPD-5 N-terminal fragment (aa 1-473) exhibited a native molecular weight consistent with being a dimer both when untagged and when fused to a monomeric MBP tag, as did an MBP-tagged version of the second largest N-terminal fragment aa 1-345 (**Fig. 5A, Fig. S3A**). Testing the larger untagged fragment in the presence and absence of PLK1 phosphorylation revealed that phosphorylation had no effect on native molecular weight (**Fig. 5A, Fig. S3A**). Thus, the SPD-5 N-terminus is a dimer, regardless of PLK1 phosphorylation status.

**Fig. 5.**
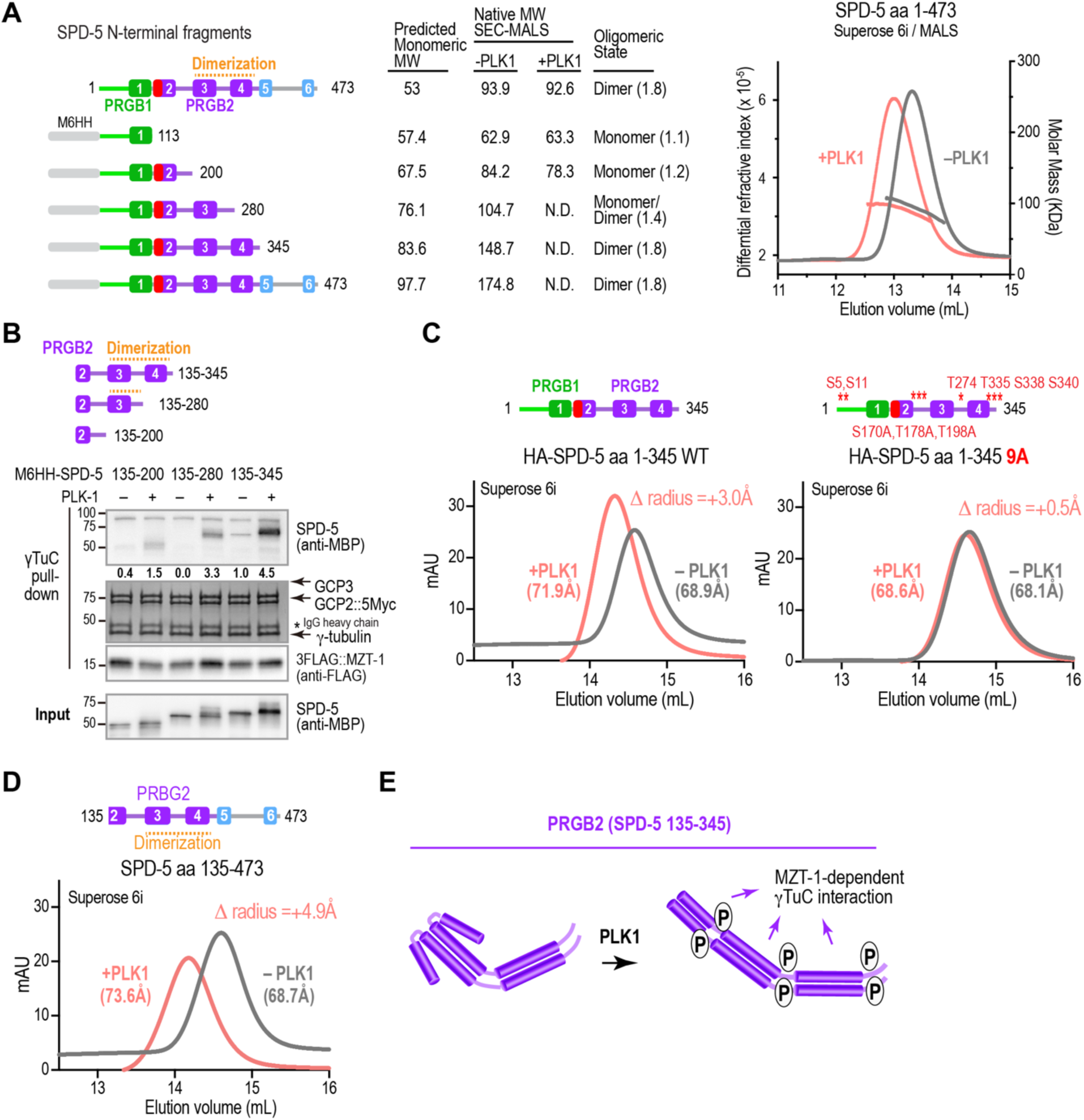
The SPD-5 N-terminus is dimerized by PRGB2 and undergoes a conformational change upon PLK1 phosphorylation. **(A)** (*left*) Schematics of untagged and MBP-6His-HA (M6HH)-tagged SPD-5 variants that were analyzed by SEC-MALS. (*middle*) Table summarizing SEC-MALS results. For each SPD-5 fragment, the Table lists the predicted monomeric molecular weight, the native molecular weight measured by SEC-MALS with or without PLK1 phosphorylation and interpreted oligomerization state with ratio of native to predicted MW in parentheses; *N.D.*: not determined. (*right*) Example SEC-MALS data for untagged SPD-5 aa 1-473 before and after PLK1 phosphorylation. While phosphorylated and unphosphorylated dimers have the same native molecular weight as measured by MALS, the phosphorylated dimers elute earlier from the column, suggesting they have an increased hydrodynamic radius. **(B)** Binding assays with γTuC-coated beads and the indicated MBP-6His-HA (M6HH)-tagged SPD-5 fragments preincubated with or without PLK1. SPD-5 and MZT-1 were analyzed by immunoblotting using the indicated antibodies; γ-tubulin, GCP2^GIP-2^, and GCP3^GIP-1^ were detected by Coomassie staining. Numbers below the SPD-5 fragment bands indicate band intensity relative to SPD-5 135-345 in the absence of PLK1 phosphorylation. Asterisk indicates the IgG heavy chain of the anti-Myc antibody used for the Myc IP. **(C)** Size exclusion chromatography analysis of HA-tagged SPD-5 aa 1-345 (*left*) or the same fragment with 9 indicated predicted PLK1 sites mutated to alanines (*right*). The hydrodynamic radii of the SPD-5 fragments were calculated based on column calibration with standard proteins and are shown in parentheses; the change in hydrodynamic radius induced by phosphorylation is shown in the upper right corner of each graph. **(D)** Size exclusion chromatography analysis of SPD-5 aa 135-473 (lacking PRGB1 and γTuNA-In) after preincubation with or without PLK1 as indicated. Hydrodynamic radii measurements are shown as in *(C)*. **(E)** Speculative model of how PLK1 phosphorylation promotes a conformational change in PRGB2 and also positively contributes affinity to promote γTuC binding.

In contrast to the two large N-terminal fragments, PRGB1 alone (SPD-5 aa 1-113) and the longer aa 1-200 fragment both had native molecular weights consistent with being monomeric, regardless of PLK1 phosphorylation status (**Fig. 5A**, **Fig. S3B**). SPD-5 aa 1-280, which contains predicted helix 3 but not 4, had a native molecular weight intermediate between monomer and dimer, suggesting a monomer-dimer equilibrium (**Fig. 5A**; **Fig. S3A**). Collectively, the SEC-MALS analysis indicates that dimerization of the SPD-5 N-terminus depends on α-helices 3 and 4 (**Fig. 5A**). Comparing γTuC binding of an MBP fusion with PRGB2 (aa 135-345), which binds robustly, to that of the same construct lacking helix 4 (aa 135-280) or helices 3 and 4 (aa 135-200) revealed almost no binding in the absence of helices 3 and 4, and weak phospho-stimulated binding when helix 3 was present and helix 4 was absent. Thus, PRGB2-mediated γTuC binding correlates with dimerization mediated by helices 3 and 4 (**Fig. 5B**).

While the native molecular weight of the full SPD-5 N-terminus (aa 1-473) was not altered by phosphorylation, the phosphorylated form eluted significantly earlier than the non-phosphorylated form on the size exclusion column (**Fig. 5A**). As size exclusion chromatography separates proteins based on hydrodynamic radius, the earlier elution suggested that phosphorylation causes the dimers to adopt a more elongated conformation. This is likely due to a conformational change in PRGB2, as a phosphorylation-dependent shift was observed for fragments containing PRGB2 (SPD-5 aa 1-345 and SPD-5 aa 135-473; **Fig. 5C-E**) but not fragments containing only PRGB1 or PRGB1 and the γTuNA-In (SPD-5 aa 1-113 or aa 1-200; **Fig. S3B**).

To confirm that the observed shift in elution on the size exclusion column was due to phosphorylation, we mutated a set of 9 predicted, high-confidence PLK1 sites in SPD-5 (aa 1-345), which contains PRGB1 followed by the γTuNA-In and PRGB2. The mutated sites included three subsets implicated in PLK1 stimulated γTuC binding *in vitro* (S5 and S11, **Fig. 1I**; S170, T178 and T198, **Fig. 3E** (*32*); and T274, T335, S339 and S340, **Fig. 3E**); the first two sets are also important for the centrosomal recruitment of γ-tubulin *in vivo*; **Fig. 2B**, (*32*)). The PLK1-dependent shift was largely abolished in the 9A mutant (**Fig. 5C**), whereas mutation of either the first 5 or last 4 of the 9 sites reduced but did not eliminate the shift in elution volume (**Fig. S3C**). These results indicate that the SPD-5 N-terminus, which has two independent γTuC-binding interfaces, is a constitutive dimer that undergoes a PLK1 phosphorylation-stimulated conformational change as the result of distributed phosphorylation across the N-terminus. Notably, a significant conformational change was still observed following mutation of the 5 N-terminal sites that are critical for γTuC binding and centrosomal recruitment (**Fig. S3C**, 5A mutant). This result suggests that, although the PLK1-stimulated conformational change is potentially required for γTuC binding, it may not be sufficient. In addition to facilitating an elongated conformation of the dimer, phosphorylation of these residues likely makes a positive contribution to γTuC-binding affinity. Collectively, these data indicate that PLK1 phosphorylation induces a conformational change of the dimeric SPD-5 N-terminus and contributes positive affinity to γTuC binding.

### PLK1 phosphorylation disrupts a potentially inhibitory interaction between PRGB1 and PRGB2

The predicted α-helix 2 that follows PRGB1 has both the γTuNA-In and a region (aa 135-180) required for γTuC-binding by PRBG2 (**Fig. 3D**). Thus, one appealing model is that interaction of α-helix 2 with the γTuNA mutually inhibits γTuC-binding by both PRGB1 and PRBG2 in the unphosphorylated state. To test this idea, we conducted binding assays between PRGB1 and fragments containing part or all of PRGB2. We found that PRGB1 (SPD-5 aa 1-101), but not SPD-5 aa 1-60 which lacks the γTuNA, bound to SPD-5 aa 101-473, which contains the γTuNA-In and PRGB2 (**Fig. 6A**). In addition, PRGB1 (aa 1-113) bound to SPD-5 aa 135-200 (the part of α-helix 2 after the γTuNA-In required for PRBG2 to bind to the γTuC) but not with aa 200-345 (**Fig. 6B**). Notably, the interaction between PRGB1 and SPD-5 aa 135-200 was disrupted by PLK1 phosphorylation. Collectively, these results suggest that PLK1 phosphorylation releases an interaction between PRGB1 and PRGB2 (**Fig. 6C**). This release is not sufficient to allow γTuC binding by N-terminal fragments containing α-helices 1 and 2 because PRGB1 is also inhibited by the γTuNA-In. As shown above, overcoming the inhibitory interaction of the γTuNA-In with the γTuNA requires interaction of PRGB2 with the MZT-1 module of the γTuC.

**Fig. 6.**
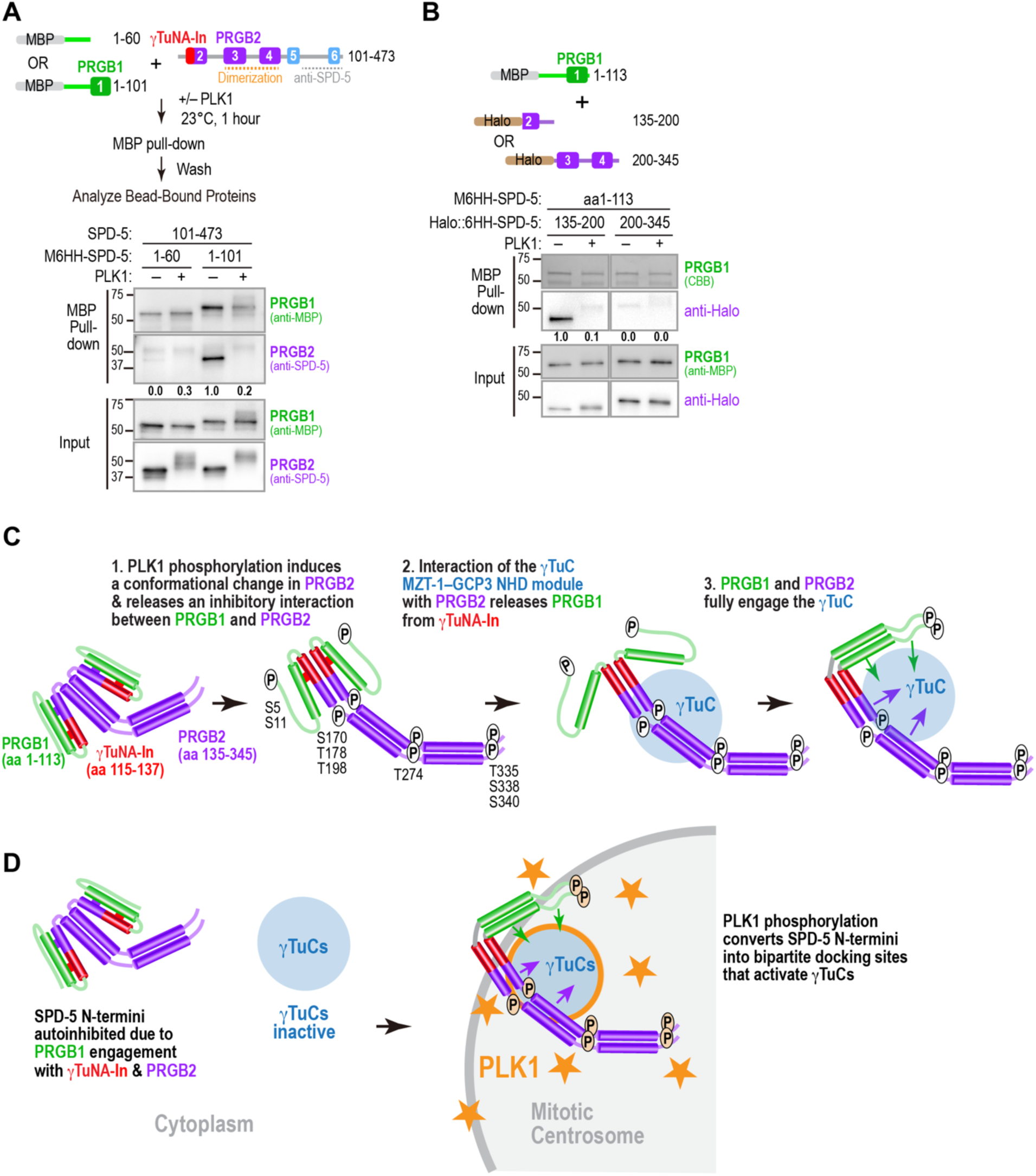
PLK1 phosphorylation disrupts an interaction between PRGB1 and PRGB2. **(A)** (*top*) Schematic of experiment to test binding between PRGB1 and SPD-5 101-473, which contains the γTuNA-In and PRGB2; (*bottom*) Results of binding analysis for the indicated conditions. MBP-6His-HA (M6HH)-tagged PRGB1 or aa1-60, which lacks the γTuNA, were mixed with untagged SPD-5 aa 101-473 containing the γTuNA-In and PRGB2. Mixtures were incubated with or without PLK1 and bead-bound proteins were detected by immunoblotting after an MBP-pull-down. Numbers below the PRGB2 indicate PRGB2 band intensity relative to the SPD-5 1-101 pull-down in the absence of PLK1 phosphorylation. **(B)** (*top*) Schematic of SPD-5 variants used to analyze binding between PRGB1 and two parts of PRGB2; (*bottom*) Results of the binding analysis; an MBP-6His-HA (M6HH)-tagged SPD-5 PRGB1 fragment was mixed with the indicated Halo-tagged SPD-5 fragments and binding was analyzed as in *(A)*. Bead-bound proteins were detected by immunoblotting using the indicated antibodies or Coomassie staining. Numbers below the Halo-tagged fragments are band intensity of the halo-tagged fragment relative to amount of SPD-5 135-200 pulled down in the absence of PLK1 phosphorylation. **(C)** Model for the phosphorylation-dependent conversion of the SPD-5 N-terminus into a γTuC docking site. PLK1 phosphorylation stimulates a conformational change that extends the dimeric PRGB2 region and releases an inhibitory interaction between PRGB1 and α-helix 2 in PRGB2. Interaction of PRGB2 with the MZT-1 module of the γTuC releases PRGB1 from inhibition by the γTuNA-In, enabling PRGB1 to engage the γTuC. Phosphorylation also likely contributes positive affinity to the interactions of PRGB1 and PRGB2 with the γTuC. **(D)** Model for the functional significance of PLK1 control of γTuC docking. The complex multi-step process required to convert the SPD-5 N-terminus into a γTuC docking site ensures that it is fully inhibited in the cytoplasm and only binds to and activates γTuCs on the centrosomal scaffold where PLK1 activity is concentrated. Tight control facilitates precise patterning of microtubule nucleation to enable rapid, robust spindle assembly.

## Discussion

During cell division in metazoans, a major function of centrosomes is to catalyze the assembly of a bipolar mitotic spindle that segregates the replicated chromosomes (*3–9*). In preparation for this role, centrosomes increase in size and microtubule nucleating capacity during mitotic entry in a process controlled by PLK1 kinase that concentrates on the PCM matrix (*10, 11, 13–16, 32, 47, 51*). PLK1 phosphorylates CDK5RAP2 family molecules to drive their assembly to form the matrix (*11, 16, 18, 19*) and to convert them into mitotic docking sites for γTuCs (*32, 34*). Here, we use the *C. elegans* CDK5RAP2 family protein SPD-5 as a model to investigate the PLK1 phosphorylation-driven generation of a mitotic γTuC docking site. Our analysis revealed that the SPD-5 N-terminus contains two regions, PRGB1 and PRGB2, that are each capable of PLK1-stimulated binding to γTuCs but which have distinct requirements on the γTuC for their binding: PRGB2 requires the MZT-1 module whereas PRBG1 does not (**Fig. 4C**). Based on our data, we propose a model for how PLK1 phosphorylation facilitates conversion of the inhibited SPD-5 N-terminus into a γTuC docking site (**Fig. 6C**). Our data suggest that PLK1 phosphorylation stimulates a conformational change that extends the dimeric PRGB2 region, releases an inhibitory interaction between the γTuNA in PRGB1 and PRGB2 that enables PRGB2 to interact with the MZT-1 module of the γTuC, which in turn releases PRGB1 from inhibition by the γTuNA-In, and contributes positive affinity to the interfaces of PRGB1 and PRGB2 with the γTuC. This complex, multi-step conversion process would ensure that neither PRGB1 nor PRGB2 interact with or activate γTuCs in the cytoplasm, thereby concentrating γTuC-mediated microtubule nucleation in the vicinity of the high PLK1 activity environment of the centrosomal scaffold in order to facilitate rapid and robust spindle assembly (**Fig. 6D**).

CDK5RAP2 family proteins contain a conserved CM1 motif (*35, 36*), which is comprised of an initial ∼30aa peptide, the γTuNA, that binds and activates γTuCs (*37–42*), followed by an autoinhibitory region, the γTuNA-In, that prevents the γTuNA from interacting with γTuCs (*44*). Work on the CDK5RAP2 family protein Cnn in *Drosophila*, has suggested that the CM1 region is also preceded by a phosphorylation-regulated autoinhibitory domain (*34*). In *C. elegans* SPD-5, the region with homology to the γTuNA (aa 59-101) is similarly flanked by inhibitory regions on both sides; inhibition by the N-terminal one (aa 1-60) is released by PLK1-phosphorylation but inhibition by the C-terminal γTuNA-In is not, and is only relieved once phosphorylated PRGB2 engages the γTuC. Given the broad conservation of the CM1 region across γTuC docking proteins (*22, 35, 44, 60*), it is likely that the bipartite functional organization of a γTuC-binding γTuNA region followed by an inhibitory γTuNA-In region is conserved.

In addition to PRGB1 and the γTuNA-In, we identified PRGB2 (SPD-5 aa 135-345), a second region capable of robust PLK1 phosphorylation-dependent γTuC binding. Thus, to prevent γTuC binding by the SPD-5 N-terminus, autoinhibitory mechanisms must prevent γTuC binding by both PRGB1 and PRGB2. Our results suggest that this may be achieved in part through mutual inhibition that is unwound in a stepwise manner by PLK1 phosphorylation. Importantly, relieving inhibition of the γTuNA by the γTuNA-In can only be accomplished in the context of dimeric PRGB2 (**Fig. 1E,G**; **Fig. 5A,B; Fig. S3A,B**). Phosphorylation triggers a conformational change broadly distributed across PRGB2 (**Fig. 3E**; **Fig. 5A,C**; **Fig. S3B,C)** that is important for binding of PRGB2 to the γTuC but also for relief of PRGB1 inhibition by γTuNA-In. Collectively, these results support a sequential model (**Fig. 6D**), where phosphorylation of the SPD-5 N-terminus by PLK1 releases the interaction between PRGB1 and PRGB2 and induces a conformational change in PRGB2, which then binds to the MZT-1–GCP3 NHD module of the γTuC. Engagement of PRGB2 with the γTuC is necessary to release inhibition of the γTuNA by the γTuNA-In and allow PRGB1 to bind. A key piece of evidence supporting this model is that the γTuC lacking the MZT-1 module fails to interact with the SPD-5 N-terminus, despite PLK1 phosphorylation.

In vertebrates, the γTuNA region of CDK5RAP2 forms a dimeric coiled-coil that interacts with MZT2–GCP2 NHD modules to form tripartite complexes that bind at the junctions between adjacent γTuSCs in pre-formed γTuRCs, linking them together to partially close the γTuRC and enhance its nucleating activity (*40–42*). In contrast, the phosphorylated PRGB1 region in SPD-5, is capable of robust γTuC interaction as a monomer, suggesting that the *C. elegans* γTuNA region does not need to be a dimeric coiled-coil to interact with γTuCs. This finding is reminiscent of prior structural work on the budding yeast pericentrin-like protein Spc110, which promotes γTuSC polymerization to form larger helical assemblies (*60, 61*). Structural work has indicated that the two CM1 regions in the Spc110 dimer contact different regions of the γTuSC, with one of the CM1 regions bridging a pair of adjacent γTuSCs as a single α-helix (*62*).

In contrast to PLK1-phosphorylated PRGB1, which binds the γTuC as a monomer independent of MZT-1, PRGB2 is dimeric and can only bind to γTuCs containing a MZT-1 module. Although the broad conservation of the CM1 region makes it seem likely that the γTuNA followed by γTuNA-In architecture is conserved (*22, 35, 44, 60*), an interesting question is whether the presence of two γTuC binding interfaces, one on either side of the inhibitory γTuNA-In, and a MZT-1-based mechanism for release of the γTuNA from the γTuNA-In will be conserved in other CM1 motif-containing proteins. We note that similar to the bipartite interface with γTuCs that we suggest here for CDK5RAP2 family proteins, work in budding yeast has suggested that the yeast γTuC receptor Spc110 and other pericentrin-related proteins contain a second motif, termed a SPM motif, positioned N-terminal to their CM1 domain, that interfaces with the γTuSC along with the CM1 region (*60*).

Collectively, the results described here suggest that the phosphorylation-based mechanisms that generate γTuC complex docking sites on the mitotic centrosome scaffold are likely to be multi-step and complex. Such complex regulation likely serves to facilitate spindle assembly by ensuring robust and precise spatiotemporal control of the generation of microtubule nucleating sites.

## Materials and Methods

### *C. elegans* strains and transgene generation

*C. elegans* strains (listed in **Table S1**) were maintained at 16°C. Single-copy transgenes were generated using the MosSCI transposon-based method (*63*) to insert them at defined chromosomal sites. Transgenes were cloned into pCFJ151, which contains the Cb-unc-119 selection marker and appropriate homology arms. SPD-5 mutant transgenes were generated as described previously (*32*). Because the endogenous *spd-5* promoter contains a highly repetitive sequence that reduces MosSCI efficiency, *spd-5* transgenes were instead driven by the *spd-2* promoter, comprising 3043 bp upstream of the start codon. The SPD-5 coding region was followed by 577 bp downstream of the stop codon and a 577 bp internal segment was re-encoded to confer resistance to a dsRNA targeting this region without altering protein coding information. Injection mixes contained the pCFJ151-derived repair plasmid (50-100 ng/μL), transposase plasmid pCFJ601 (Peft-3::Mos1 transposase, 50 ng/µl), and four plasmids for negative selection against chromosomal arrays: pMA122 (Phsp-16.41::peel-1, 10 ng/μL), pCFJ90 (Pmyo-2::mCherry, 2.5 ng/μL), pCFJ104 (Pmyo-3::mCherry, 5 ng/μL), and pGH8 (Prab-3::mCherry, 10 ng/μL). Mixes were injected into strains EG6429 or EG6699 to target the ttTi5605 site on Chr II. After one week, the progeny of injected worms were heat-shocked at 34°C for 3 hours to induce the expression of PEEL-1 to kill worms containing extra chromosomal arrays. Moving worms lacking fluorescent markers were identified as candidates, and PCR across both integration junctions was used to confirm transgene integration in their progeny.

### RNA interference

Single-stranded RNAs (ssRNAs) were synthesized in 50 μL T3 and T7 transcription reactions (MEGAscript, Invitrogen) using gel purified DNA templates generated by PCR from N2 genomic DNA with primers containing T3 or T7 promoter sequences (**Table S2**). Transcription products were purified using the MEGAclear kit (Invitrogen). Equal volumes (50 μL) of T3- and T7-generated ssRNA were combined with 50 μL of 3x soaking buffer (32.7 mM Na_2_HPO_4_, 16.5 mM KH_2_PO_4_, 6.3 mM NaCl, 14.1 mM NH_4_Cl) and annealed by heating at 68°C for 10 minutes, followed by 37°C for 30 minutes to form double-stranded RNA (dsRNA). dsRNAs were injected at a concentration of ≥ 1.3 μg/μL. For live imaging of early embryos after RNAi, L4 hermaphrodites were injected with dsRNAs and incubated at 16°C for 48 hours before dissection to isolate embryos for imaging.

### Antibodies

A previously described antibody against SPD-5 (392-550 aa) (*64*) was used at 1 μg/mL for immunoblotting. The following commercially available antibodies were used at the indicated dilutions: anti-α-tubulin (DM1A; Sigma-Aldrich; 1:5,000), anti-FLAG (F1804; Sigma-Aldrich; 1:1,000), anti-Myc (9E10; M4439; Sigma-Aldrich; 1:5,000), anti-MBP (ab9084; Abcam; 1:1,000), anti-MBP (E8032S; NEB; 1:5,000), anti-Halo (G9211; Promega; 1:1,000). Secondary antibodies were obtained from Jackson ImmunoResearch and GE Healthcare.

### Live imaging

Embryos for live imaging experiments were obtained by dissecting gravid adult hermaphrodites in M9 buffer (42 mM Na_2_HPO_4_, 22 mM KH_2_PO_4_, 86 mM NaCl, and 1 mM MgSO_4_). One-cell embryos were transferred with a mouth pipette onto a 2% agarose pad and overlaid with a 22 x 22-mm coverslip. Imaging was performed in a temperature-controlled room at 20°C using spinning-disk or laser-scanning confocal microscopy. Spinning-disk imaging was conducted on either (1) a Zeiss Axio Observer.Z1 inverted microscope equipped with a CSU-X1 (Yokogawa), 63x/1.4NA Plan-Apochromat objective, and a Photometrics QuantEM: 512SC camera (**Fig. 2B**), or (2) an inverted Nikon Eclipse Ti2 microscope equipped with a CSU-X1 (Yokogawa), a 60x/1.4NA Plan-Apochromat objective (Nikon), and an iXon Life EMCCD camera (iXON-L-888; Andor) (**Fig**. **2D**). Embryos shown in Figs. 2A, 2C, and 3B were imaged using a Zeiss LSM880 confocal microscope equipped with Airyscan and a 63x/1.40 Oil Plan-Apochromat WD0.19 lens (Zeiss) in a 20-22°C room. Images were processed with ZEN 2.3 SP1 Black software (Zeiss). Images of centrosomal fluorescence were acquired every 30 seconds by collecting 11 z-planes at 1.5 µm intervals without binning. Imaging was initiated in one-cell embryos between centrosome separation and pronuclear meeting and was terminated after initiation of cytokinesis.

### Image analysis

All images were processed and analyzed using ImageJ (National Institutes of Health). Centrosomal fluorescence was quantified from maximum intensity projections of entire z-stacks. To quantify centrosomal fluorescence, a fixed size box was drawn around the centrosome at each time point (smallest box that could enclose the centrosomal signal at their largest point in the image sequence; box size varied depending on marker and imaging conditions), along with a box one pixel larger on each side in both dimensions. The per-pixel background was calculated as [(integrated intensity in the larger box - integrated intensity in the smaller box)/(area of larger box - area of smaller box)]. The centrosomal signal was calculated as the mean intensity in the smaller box minus the per-pixel background. To quantify immunoblot band intensity, a fixed size box was drawn around each band (the smallest box that could enclose the strongest band signal in the blot; box size varied depending on the blot and imaging conditions), along with a box one pixel larger above and below. The per-pixel background was calculated as [(integrated intensity in the larger box - integrated intensity in the smaller box)/(area of larger box - area of smaller box)]. The band intensity was calculated as the mean intensity in the smaller box minus the per-pixel background. For γTuC pull-down assays, the SPD-5 band intensity was normalized by dividing by the corresponding band intensity in the input samples.

### Protein expression and purification

GST-, MBP-, or Halo-tagged fragments of the SPD-5 N terminus were expressed in BL21(DE3) pLysS *E. coli* from DNA constructs cloned into a pGEX-6P-1, pMAL-6His-TEV, or pHalo-6His-TEV vectors, respectively. The pMH-Halo tag plasmid was a gift from Michael Huen (Addgene plasmid #154144; http://n2t.net/addgene:154144). When bacterial cultures reached an OD_600_ of 0.6, protein expression was induced for 16-18 h at 15°C by adding 0.3 mM IPTG. Cells were washed once with cold PBS and flash frozen in liquid nitrogen. For GST-tagged proteins, pelleted cells were resuspended in lysis buffer (PBS containing 250 mM NaCl, 10 mM EGTA, 10 mM EDTA, 0.1% Tween, 200 μg/mL lysozyme, 2 mM benzamidine, and EDTA-free protease inhibitor cocktail [Roche]). For MBP-6His- or Halo-6His-tagged proteins, cells were resuspended in the same buffer supplemented with 10 mM imidazole. Cell lysis was performed by sonication, and lysates were clarified by centrifugation at 40,000 rpm for 20 minutes at 4°C in a 45 Ti rotor (Beckman). Cleared cell lysates were incubated with 500 µl of glutathione agarose (Cytiva) or Ni-NTA agarose beads (Qiagen) for 2 hours at 4°C. The resin was washed twice with 30 mL of wash buffer (PBS containing 250 mM NaCl, 1 mM β-mercaptoethanol, and 2 mM benzamidine) for GST-tagged proteins or with the same wash buffer supplemented with 20 mM Imidazole for MBP-6His- or Halo-6His-tagged proteins, followed by incubation with 10 mL washing buffer containing 5 mM ATP for 10 min at 4°C to reduce non-specific interactions with heatshock proteins. The resin was then washed three additional times with wash buffer. For GST-tagged proteins, the resin was incubated overnight at 4°C with PreScission protease (Eton Bioscience) or HRV-3C protease (ACRO Biosystems) in elution buffer (20 mM Tris-Cl pH 8.0, 150 mM NaCl and 1 mM DTT) to elute the untagged SPD-5 N-terminal fragments by cleavage from the GST-tag. The supernatant was collected the next day and stored at 4°C. For MBP-6His- or Halo-6His-tagged proteins, the resin was transferred to a poly-prep chromatography column (Bio-Rad), and eluted with buffer containing 250 mM imidazole, then stored at 4°C. The eluted 6His-tagged proteins were concentrated using 10 KDa MWCO Amicon Ultra concentrators (Millipore) and dialyzed into 20 mM Tris-HCl (pH 7.5), 50 mM NaCl, 2 mM MgCl_2_, and 1 mM DTT. C-terminally 6His-tagged PLK1 T194D, purified from Sf9 cells, was a gift from Jeffrey Woodruff (UT Southwestern).

### Purification of reconstituted *C. elegans* γ-tubulin complexes

For reconstitution of the *C. elegans* γ-tubulin complex, Myc-tagged GCP2^GIP-2^, FLAG-tagged MZT-1, GCP3^GIP-1^, and γ-tubulin^TBG-1^ were cloned into CMV-driven human expression vectors (**Table S3**) and co-transfected into FreeStyle 293-F cells (Thermo Fisher Scientific). The coding sequences for GCP2^GIP-2^ and γ-tubulin^TBG-1^ were amplified by PCR from an N2 cDNA library, whereas codon-optimized GCP3^GIP-1^ and MZT-1 sequences were synthesized (GENEWIZ). Empty 5Myc plasmid (CS2P, Addgene #17095) or 3FLAG plasmid (p3XFLAG-CMVTM-7.1, Sigma #E7533) served as negative controls. Cell transfection was performed using FreeStyle MAX Reagent and OptiPRO SFM according to the manufacturer’s instructions (Thermo Fisher Scientific). A total of 12.5 µg plasmid DNA was transfected into 10 mL of cells at 1 x 10^6^ cells/mL. After 43-38h, cells were harvested, washed with PBS, and resuspended in lysis buffer (20 mM Tris/HCl pH 7.5, 50 mM NaCl, 0.5% Triton X-100, 5 mM EGTA, 1 mM dithiothreitol (DTT), 2 mM MgCl_2_ and EDTA-free protease inhibitor cocktail [Roche]). Lysis was performed in an ice-cold sonicating water bath for 5 minutes, followed by centrifugation at 15,000x g for 15 minutes at 4 °C. Cleared lysates were incubated with Pierce Anti-c-Myc magnetic beads (Thermo Fisher Scientific) for 2 hours at 4 °C. Beads were washed five times with lysis buffer and either used directly for pull-down assay or resuspended in SDS sample buffer. For immunoblotting, equal volumes of samples were run on Mini-PROTEAN gels (Bio-Rad) and transferred to PVDF membranes using a TransBlot Turbo system (Bio-Rad). Blocking and antibody incubations were performed in TBS-T plus 5% nonfat dry milk or in TBS-T plus 5% BSA. Membranes were incubated with primary antibodies overnight at 4°C and with the secondary antibodies for 1 hour at room temperature. Signals were developed using SuperSignal West Femto substarte (Thermo Fisher Scientific) and detected using a ChemiDoc XRS+ system (BioRad). For Coomassie staining to detect *C. elegans* γ-tubulin complex, equal volumes of samples were run on Mini-PROTEAN gels (Bio-Rad) and stained with SimplyBlue^TM^ Safe Stain (Thermo Fisher).

### Kinase and pulldown assays

For the γ-tubulin complex pulldown assays (Figs. 1, 3-5), each purified SPD-5 fragment (final concentration 2.0 μM) of was incubated with constitutively active PLK1(T194D) (500 nM final concentration) in kinase buffer (20 mM Tris-Cl pH 7.5, 50 mM NaCl, 10 mM MgCl_2_, 0.2 mM ATP, 1 mM DTT) for 1 hour at 23°C. Following phosphorylation, the reaction mixtures were combined with γTuCs immobilized on Myc beads in lysis buffer and incubated for 2 hours at 4°C. The final concentration of the SPD-5 fragments was 75 nM. Beads were washed five times with lysis buffer and resuspended in sample buffer prior to analysis on SDS-PAGE. For the MBP pull-down assays (Fig. 6), purified proteins were mixed at a final concentration of 2.0 μM with 500 nM constitutively active PLK1(T194D) in kinase buffer for 1 hour at 23°C. Reaction mixtures were then mixed with 2 µg of anti-MBP antibodies (Abcam) in lysis buffer and incubated for 2 hours at 4°C. The final concentration of each purified SPD-5 fragment was 75 nM. After the incubation, the proteins were incubated with protein A beads (Thermo Fisher, #88845) for 1 hour at 4°C. The beads were washed five times with lysis buffer and resuspended in sample buffer prior to analysis on SDS-PAGE.

### Gel filtration assay

Purified proteins were concentrated as necessary using 10 KDa MWCO Amicon Ultra concentrators (Millipore) and subjected to size-exclusion chromatography on a Superose 6 increase 10/300 or a Superdex 200 increase 10/300 gel filtration column (Cytiva) equilibrated in gel filtration buffer (20 mM Tris-HCl (pH 7.5), 50 mM NaCl, 5 mM EGTA, 2 mM MgCl_2_, and 1 mM DTT). Peak fractions were collected, reconcentrated using 10 KDa MWCO Amicon Ultra concentrators (Millipore), and analyzed by SDS-Page and Coomassie staining. To assess whether there was a shift in native molecular weight or hydrodynamic radius of SPD-5 N fragments after PLK1 treatment, 25 µM of purified proteins by gel filtration were mixed with or without 1 µM of constitutive active PLK1-6His in the gel filtration buffer containing 5 mM MgCl_2_ and 0.2 mM ATP, and incubated for 1 hour at 23 °C. After the incubation, the reaction mixture was subjected to a size-exclusion chromatography on Superose 6 increase 10/300 (Cytiva) gel filtration column to analyze their hydrodynamic radius. To estimate hydrodynamic radius, 2 mg/mL of Thyroglobulin (Stroke’s radius: 85.0 Å), 0.3 mg/mL of Ferritin (61.0 Å), 3 mg/mL of Aldolase (48.1 Å), and 4 mg/mL of Ovalbumin (30.5 Å) were used as standards.

### Multiangle light scattering analysis (MALS)

Prior to analysis of purified proteins, a Superose 6 increase 10/300 or Superdex 200 increase 10/300 gel filtration column (Cytiva) was equilibrated with gel filtration buffer overnight at room temperature. 120 µL of each purified protein (∼2.0 mg/ml) was auto-loaded onto the column. Elution and light scattering of the tested proteins were monitored using Wyatt Technology HPLC software. After each run, baselines of UV absorbance and light scattering were established and protein peaks were defined to obtain the predicted molecular weights. SEC-MALS data was exported and graphed using GraphPad Prism software.

## Funding

National Institutes of Health grant R01GM074207 (KO)

National Institutes of Health grant R01GM074215 (AD)

National Institutes of Health grant R35 GM144121 (KDC)

Japan Society for the Promotion of Science KAKENHI 24K09461 (MO)

Buribushi fellowship from the Okinawa Institute of Science (MO)

Japan Science and Technology Agency FOREST program JPMJFR2432 (MO)

Partial salary support from the Ludwig Institute for Cancer Research (KO)

## Author contributions

Conceptualization: MO, OA, YG, KDC, AD, KO

Methodology: MO, OA, YG, KDC, AD,

KO Formal analysis: MO, OA, YG

Investigation: MO, OA, YG, WT

Resources: MO, KDC, AD, KO

Data curation: MO, OA, YG

Writing—original draft: MO, AD, KO

Writing—review & editing: MO, OA, YG, KDC, AD, KO

Visualization: MO, OA, YG, AD, KO

Supervision: MO, KDC, AD, KO

Project administration: MO, KDC, AD, KO

Funding acquisition: MO, KDC, AD, KO

## Competing interests

Authors declare that they have no competing interests.

## Data and materials availability

All data are available in the main text or the supplementary materials.

## Supplementary Figures and Tables

**Fig. S1.**
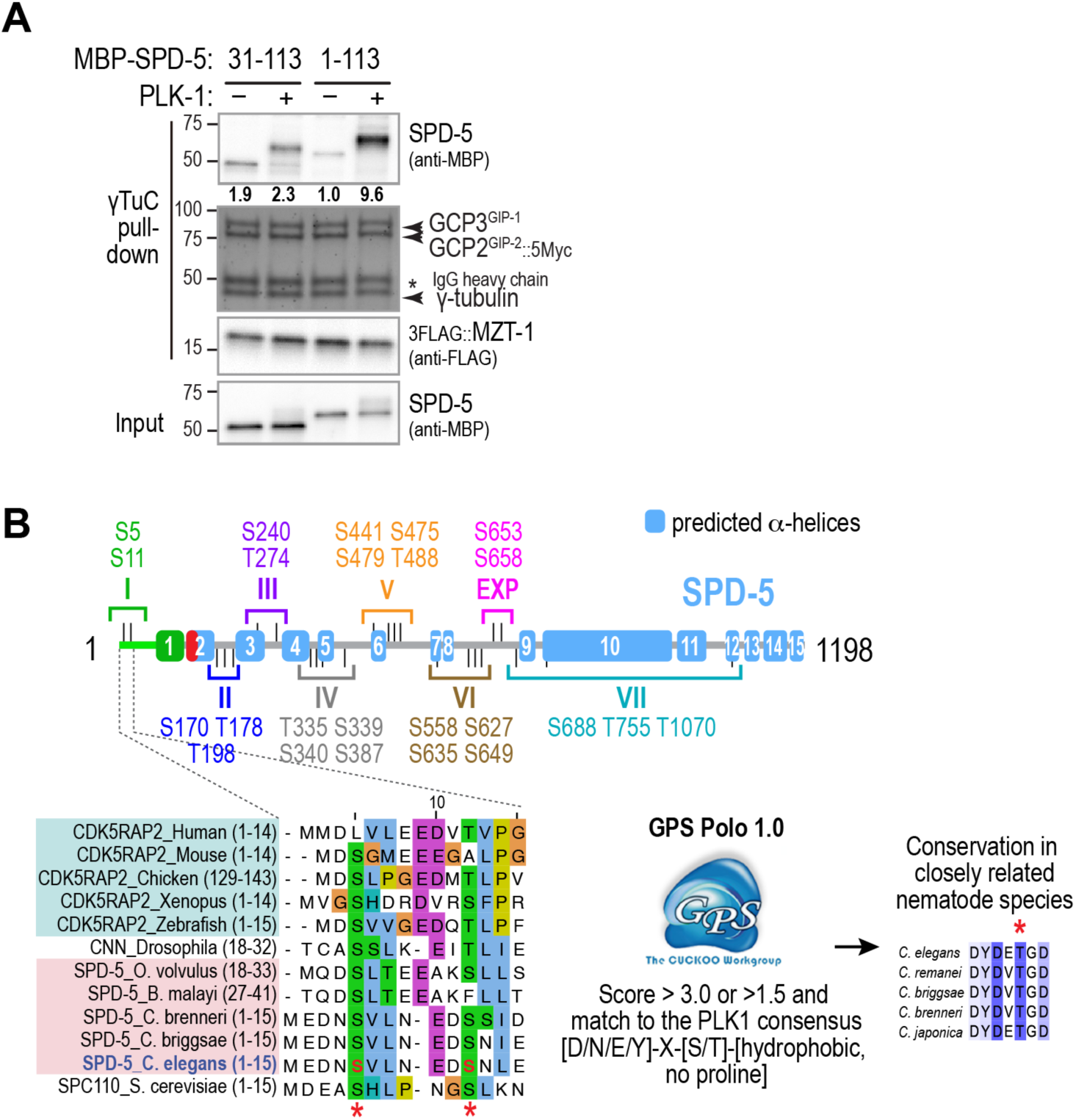
Putative PLK1 target sites in PRGB1 and their sequence alignment. (**A**) Binding assays, conducted as outlined in Fig. 1C, with γTuC-coated beads and MBP-6His-HA (M6HH)-tagged SPD-5 fragments, preincubated with or without PLK1 as indicated. SPD-5 and MZT-1 were analyzed by immunoblotting using the indicated antibodies; γ-tubulin, GCP2^GIP-2^, and GCP3^GIP-1^ were detected by Coomassie staining. Numbers below the SPD-5 fragment bands indicate band intensity relative to SPD-5 1-113 in the absence of PLK1 phosphorylation. Asterisk indicates the IgG heavy chain of the anti-Myc antibody used for the Myc IP. The blot for SPD-5 1-113 is the same as that used in Fig. 1E. (**B**) Top: Schematic depicting candidate PLK1 sites identified using the method shown in the lower right that were mutated in the indicated regional clusters in a prior study that identified cluster II as causing penetrant embryonic lethality (*32*). Bottom Left: Alignment of the indicated sequences from the N-termini of CDK5RAP2 family proteins across species, highlighting potential conservation of the S5 and S11 sites. Phosphorylation sites in alignments are marked with red asterisks.

**Fig. S2.**
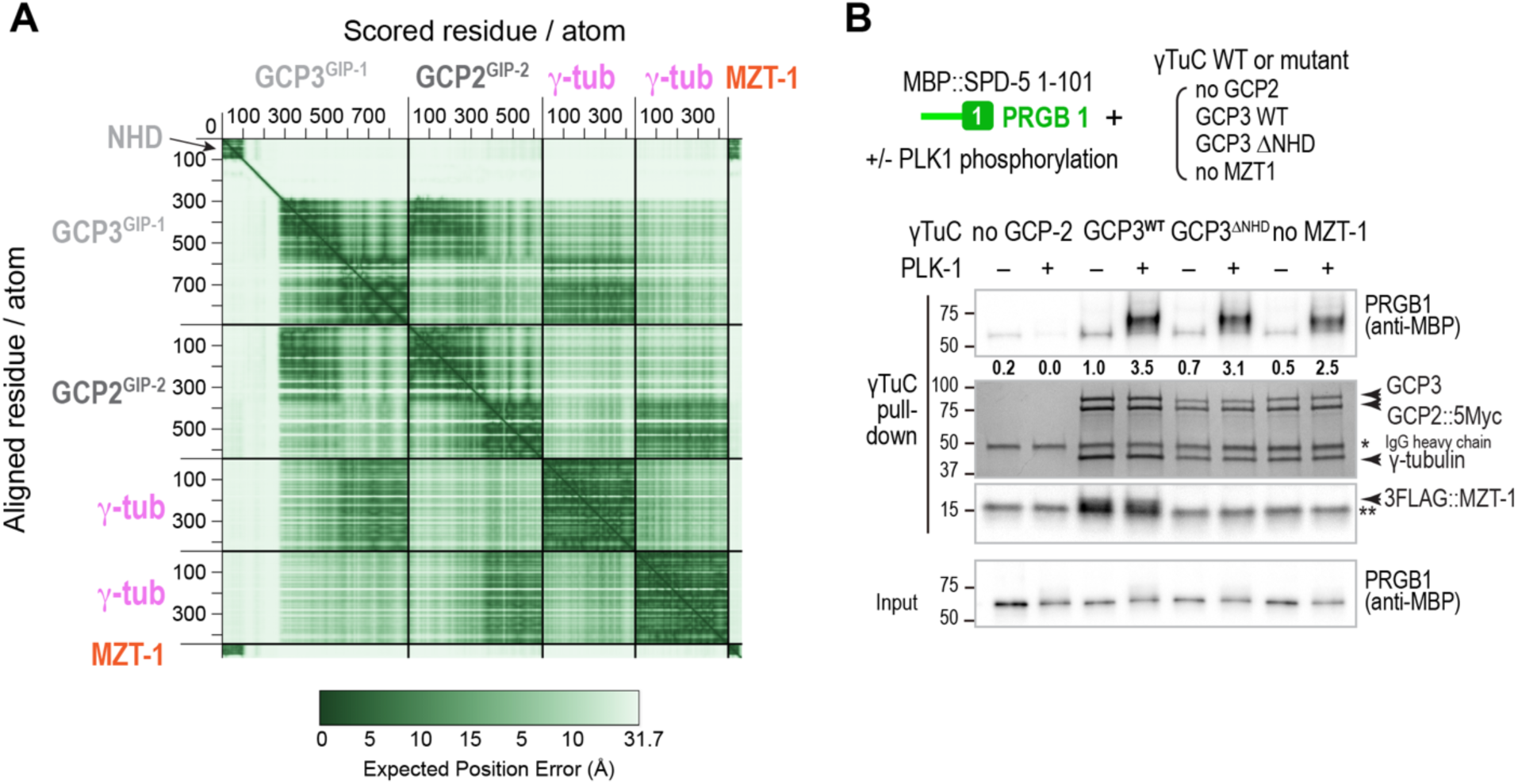
MZT-1 is dispensable for PRGB1 binding to γTuC. **(A)** Predicted Aligned Error (PAE) plot generated using PAEViewer (*66*) for the *C. elegans* γTuC model in Fig. 4A. The MZT-1–GCP3^GIP-1^ NHD module is separated by a disordered linker and is not positioned relative to the core heterotetrameric complex. **(B)** Expanded version of the western blots/coomassie gel shown in Fig. 4C that includes controls in which the plasmids encoding GCP2 which has the Myc tag for the IP (*left two lanes*) or MZT-1 (right two lanes) were omitted during γTuC assembly. Like deletion of the GCP3^GIP-1^ NHD, assembling the γTuC without MZT-1 also does not impact its PLK1-dependent binding to PRGB1. Beads prepared in with the indicated γTuC components were incubated with MBP-6His-HA-tagged SPD-5 aa 1-101 after preincubation with or without PLK1 as indicated. SPD-5 and MZT-1 were analyzed by immunoblotting using the indicated antibodies; γ-tubulin, GCP2^GIP-2^, and GCP3^GIP-1^ were detected by Coomassie staining. Numbers below the PRGB1 bands indicate band intensity relative to PRGB1 pulled down by γTuC assembled in the presence of WT GCP3 in the absence of PLK1 phosphorylation. Single asterisk indicates the IgG heavy chain of the anti-Myc antibody used for the Myc IP. Double asterisk marks the location of a non-specific band.

**Fig. S3.**
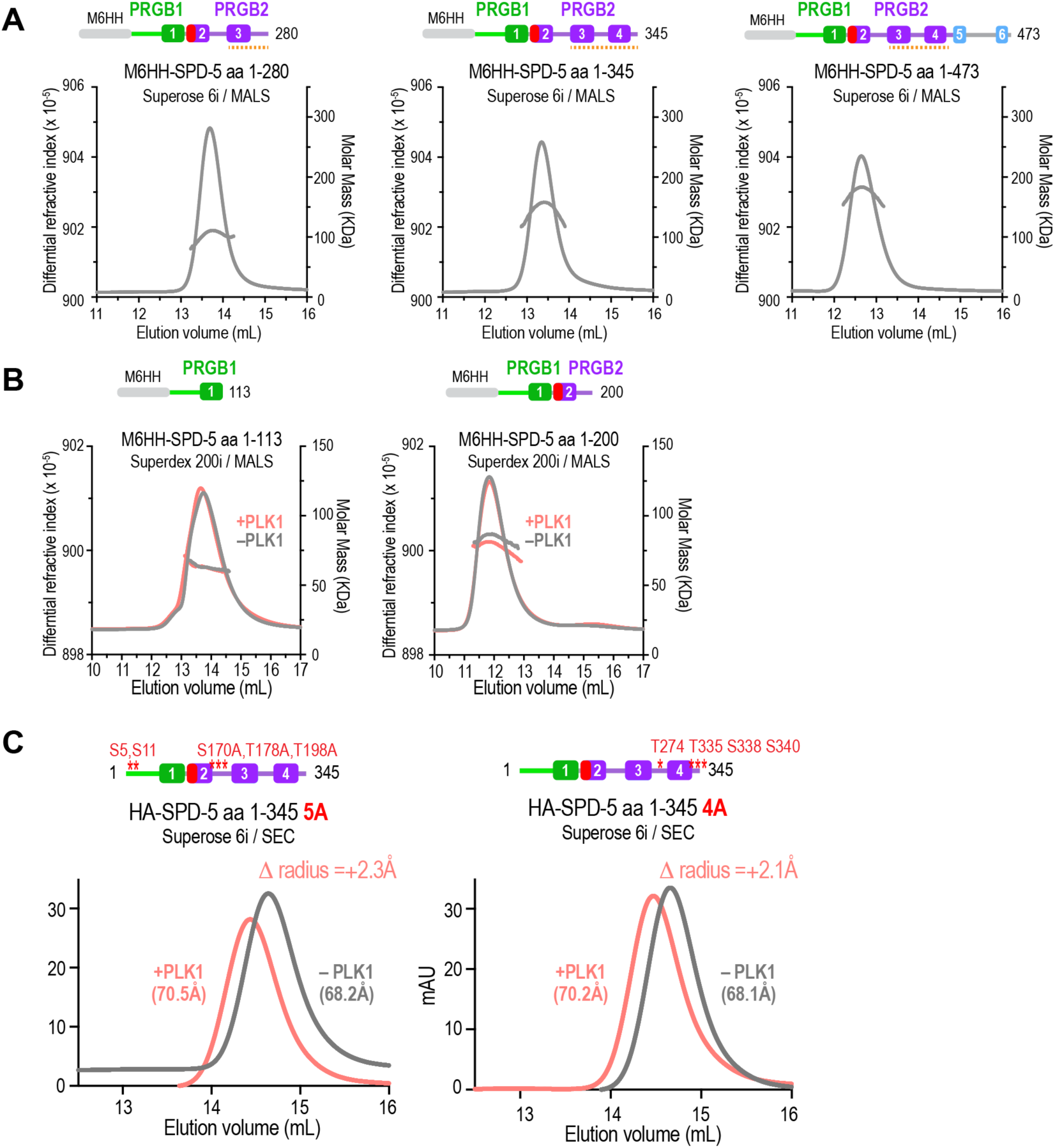
The SPD-5 N-terminus undergoes a PLK1-stimulated conformational change as the result of distributed phosphorylation across the N-terminus. **(A)** SEC-MALS data for MBP-6His-HA-tagged (M6HH)-tagged SPD-5 aa 1-280 and SPD-5 aa 1-345 used to determine the values reported in Fig. 5A. **(B)** SEC-MALS data for MBP-6His-HA-tagged (M6HH)-tagged SPD-5 aa 1-113 and SPD-5 aa 1-200 before and after PLK1 phosphorylation used to determine the values in Fig. 5A. The two fragments are monomeric both in the presence and absence of PLK1 phosphorylation and do not exhibit a phosphorylation-dependent shift in elution volume. Orange dashed line marks the region required for dimerization. **(C)** Size exclusion chromatography (SEC) analysis of HA-tagged SPD-5 aa 1-345 with the indicated sets of predicted PLK1 sites mutated to alanine after preincubation with or without PLK1 as indicated. The hydrodynamic radii of the SPD-5 fragments were calculated based on standard proteins and are shown in parentheses and the change in radius induced by phosphorylation is in the upper right corner of each graph.

**Table S1.** *C. elegans* strains used in this study.

| Strain # | Genotype | Figure |
| --- | --- | --- |
| OD4412 | ltSi569[oxTi185; pOD1110/pSW008; CEOP3608 TBG-1::mCherry; cb-unc-119(+)]I; ltSi1216 [pOD1021/pVV103; Pspd-2::GFP::SPD-5 reencoded; cb-unc-119(+)]II; unc-119(ed3) III | 2A-C, 3B, |
| MOW11 | ltSi569[oxTi185; pOD1110/pSW008; CEOP3608 TBG-1::mCherry; cb-unc-119(+)]I; ltSi1967 [pMO1326; Pspd-2::GFP::SPD-5 reencoded (deletion aa61-93)::spd-5 3'UTR; cb-unc-119(+)]II | 2A |
| OD5196 | ltSi569[oxTi185; pOD1110/pSW008; CEOP3608 TBG-1::mCherry; cb-unc-119(+)]I; ltSi1532 [pMO91; Pspd-2::gfp::spd-5 S5A/S11A::spd-5 3'UTR; cb-unc-119(+)]II; unc-119(ed3) III | 2B |
| OD5220 | ltSi569[oxTi185; pOD1110/pSW008; CEOP3608 TBG-1::mCherry; cb-unc-119(+)]I; ltSi1647 [pMO253; Pspd-2::GFP::SPD-5 aa61-1198 reencoded; cb-unc-119(+)]II; unc-119(ed3) III | 2B |
| MOW09 | ltSi569[oxTi185; pOD1110/pSW008; CEOP3608 TBG-1::mCherry; cb-unc-119(+)]I; ltSi1704 [pMO269; Pspd-2::GFP::SPD-5 aa264-1198 reencoded; cb-unc-119(+)]II; unc-119(ed3) III | 2C |
| MOW21 | ltSi569[oxTi185; pOD1110/pSW008; CEOP3608 TBG-1::mCherry; cb-unc-119(+)]I; ltSi2010 [pMO1503; Pspd-2::GFP::SPD-5 reencoded (deletion aa135-180)::spd-5 3'UTR; cb-unc-119(+)]II; unc-119(ed3)III (#18) | 3B |
| OD4211 | ltSi1216[pOD1021/pVV103; Pspd-2::GFP::SPD-5 reencoded; cb-unc-119(+)]II; unc-119(ed3) III | 2D |
| OD5160 | ltSi1641 [pMO252; Pspd-2::GFP::SPD-5 aa114-1198 reencoded; cb-unc-119(+)]II; unc-119(ed3) III | 2D |
| OD5262 | ltSi1704[pMO269; Pspd-2::GFP::SPD-5 aa264-1198 reencoded; cb-unc-119(+)]II; unc-119(ed3) III | 2D |

**Table S2.**
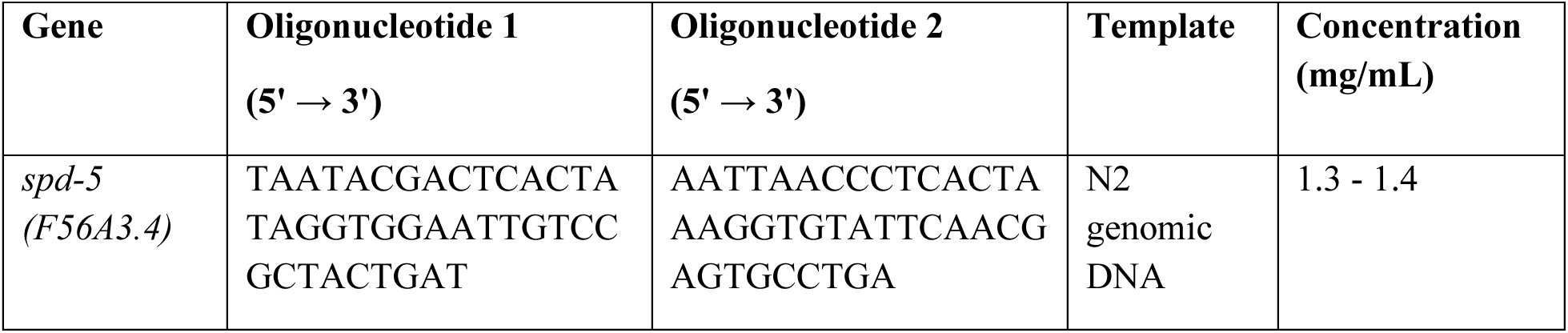
Oligos used for dsRNA production.

**Table S3.**
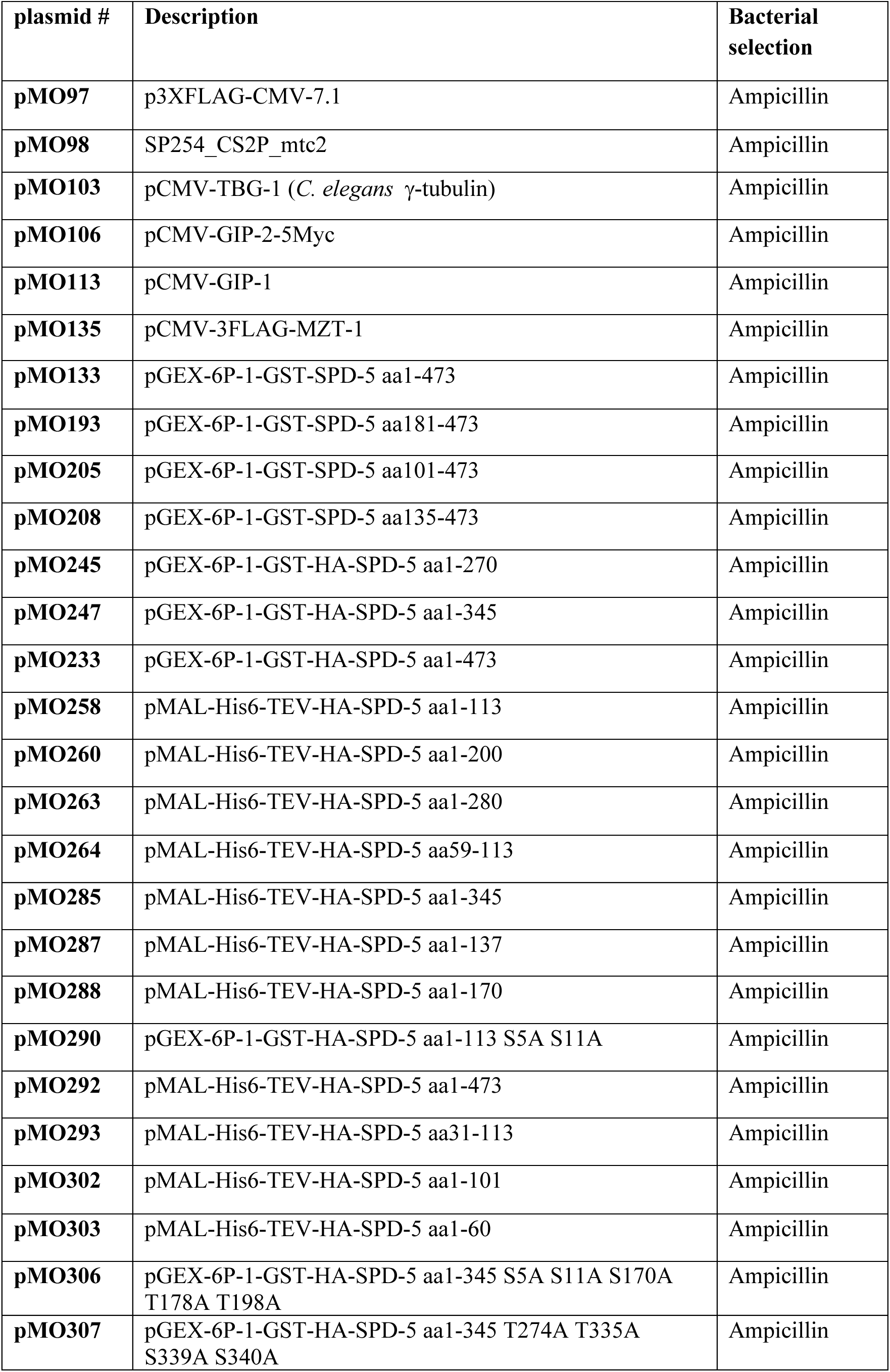
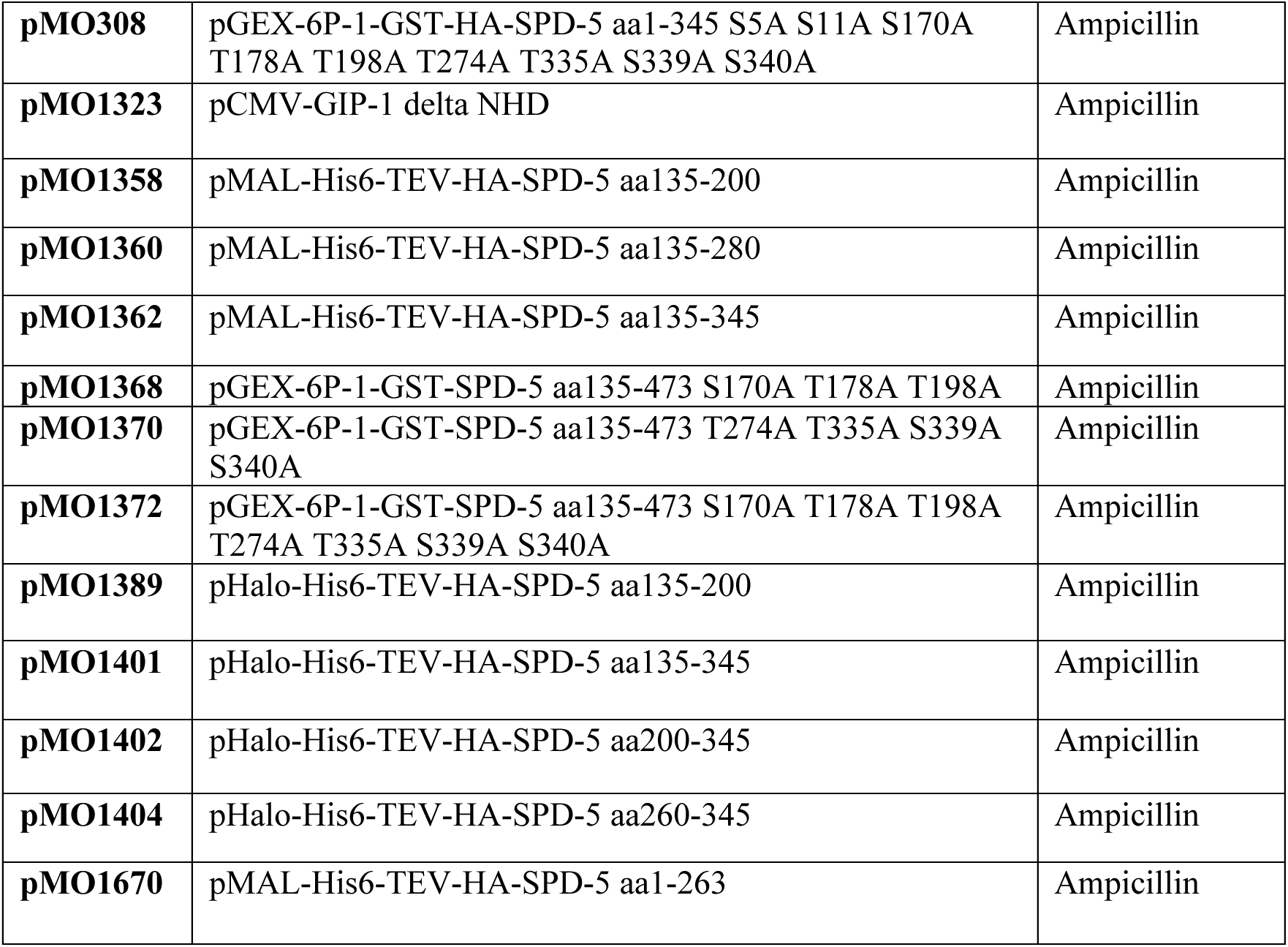
Plasmids used in this study.

## Notes

### Competing Interest Statement

The authors have declared no competing interest.

## References

1. P. T. Conduit, A. Wainman, J. W. Raff, Centrosome function and assembly in animal cells. Nat Rev Mol Cell Biol 16, 611–624 (2015).

2. S. Gomes Pereira, M. A. Dias Louro, M. Bettencourt-Dias, Biophysical and Quantitative Principles of Centrosome Biogenesis and Structure. Annu Rev Cell Dev Biol 37, 43–63 (2021).

3. H. Bazzi, K. V. Anderson, Acentriolar mitosis activates a p53-dependent apoptosis pathway in the mouse embryo. Proc Natl Acad Sci U S A 111, E1491–1500 (2014).

4. T. M. Kapoor, Metaphase Spindle Assembly. Biology (Basel) 6, (2017).

5. A. Khodjakov, C. L. Rieder, Centrosomes enhance the fidelity of cytokinesis in vertebrates and are required for cell cycle progression. J Cell Biol 153, 237–242 (2001).

6. F. Meitinger, J. V. Anzola, M. Kaulich, A. Richardson, J. D. Stender, C. Benner, C. K. Glass, S. F. Dowdy, A. Desai, A. K. Shiau, K. Oegema, 53BP1 and USP28 mediate p53 activation and G1 arrest after centrosome loss or extended mitotic duration. J Cell Biol 214, 155–166 (2016).

7. L. Pintard, B. Bowerman, Mitotic Cell Division in Caenorhabditis elegans. Genetics 211, 35–73 (2019).

8. J. H. Sir, M. Putz, O. Daly, C. G. Morrison, M. Dunning, J. V. Kilmartin, F. Gergely, Loss of centrioles causes chromosomal instability in vertebrate somatic cells. J Cell Biol 203, 747–756 (2013).

9. Y. L. Wong, J. V. Anzola, R. L. Davis, M. Yoon, A. Motamedi, A. Kroll, C. P. Seo, J. E. Hsia, S. K. Kim, J. W. Mitchell, B. J. Mitchell, A. Desai, T. C. Gahman, A. K. Shiau, K. Oegema, Cell biology. Reversible centriole depletion with an inhibitor of Polo-like kinase 4. Science 348, 1155– 1160 (2015).

10. G. Cabral, T. Laos, J. Dumont, A. Dammermann, Differential Requirements for Centrioles in Mitotic Centrosome Growth and Maintenance. Dev Cell 50, 355–366 e356 (2019).

11. P. T. Conduit, Z. Feng, J. H. Richens, J. Baumbach, A. Wainman, S. D. Bakshi, J. Dobbelaere, S. Johnson, S. M. Lea, J. W. Raff, The centrosome-specific phosphorylation of Cnn by Polo/Plk1 drives Cnn scaffold assembly and centrosome maturation. Dev Cell 28, 659–669 (2014).

12. J. Dobbelaere, F. Josue, S. Suijkerbuijk, B. Baum, N. Tapon, J. Raff, A genome-wide RNAi screen to dissect centriole duplication and centrosome maturation in Drosophila. PLoS Biol 6, e224 (2008).

13. L. Haren, T. Stearns, J. Luders, Plk1-dependent recruitment of gamma-tubulin complexes to mitotic centrosomes involves multiple PCM components. PLoS One 4, e5976 (2009).

14. H. A. Lane, E. A. Nigg, Antibody microinjection reveals an essential role for human polo-like kinase 1 (Plk1) in the functional maturation of mitotic centrosomes. J Cell Biol 135, 1701–1713 (1996).

15. K. Lee, K. Rhee, PLK1 phosphorylation of pericentrin initiates centrosome maturation at the onset of mitosis. J Cell Biol 195, 1093–1101 (2011).

16. J. B. Woodruff, O. Wueseke, V. Viscardi, J. Mahamid, S. D. Ochoa, J. Bunkenborg, P. O. Widlund, A. Pozniakovsky, E. Zanin, S. Bahmanyar, A. Zinke, S. H. Hong, M. Decker, W. Baumeister, J. S. Andersen, K. Oegema, A. A. Hyman, Centrosomes. Regulated assembly of a supramolecular centrosome scaffold in vitro. Science 348, 808–812 (2015).

17. J. B. Woodruff, O. Wueseke, A. A. Hyman, Pericentriolar material structure and dynamics. Philos Trans R Soc Lond B Biol Sci 369, (2014).

18. Y. R. Citron, C. J. Fagerstrom, B. Keszthelyi, B. Huang, N. M. Rusan, M. J. S. Kelly, D. A. Agard, The centrosomin CM2 domain is a multi-functional binding domain with distinct cell cycle roles. PLoS One 13, e0190530 (2018).

19. Z. Feng, A. Caballe, A. Wainman, S. Johnson, A. F. M. Haensele, M. A. Cottee, P. T. Conduit, S. M. Lea, J. W. Raff, Structural Basis for Mitotic Centrosome Assembly in Flies. Cell 169, 1078–1089 e1013 (2017).

20. J. M. Kollman, A. Merdes, L. Mourey, D. A. Agard, Microtubule nucleation by gamma-tubulin complexes. Nat Rev Mol Cell Biol 12, 709–721 (2011).

21. B. R. Oakley, gamma-Tubulin. Curr Top Dev Biol 49, 27–54 (2000).

22. C. A. Tovey, P. T. Conduit, Microtubule nucleation by gamma-tubulin complexes and beyond. Essays Biochem 62, 765–780 (2018).

23. Z. Zhu, I. Becam, C. A. Tovey, A. Elfarkouchi, E. C. Yen, F. Bernard, A. Guichet, P. T. Conduit, Multifaceted modes of gamma-tubulin complex recruitment and microtubule nucleation at mitotic centrosomes. J Cell Biol 222, (2023).

24. G. Goshima, R. Wollman, S. S. Goodwin, N. Zhang, J. M. Scholey, R. D. Vale, N. Stuurman, Genes required for mitotic spindle assembly in Drosophila S2 cells. Science 316, 417–421 (2007).

25. C. Verollet, N. Colombie, T. Daubon, H. M. Bourbon, M. Wright, B. Raynaud-Messina, Drosophila melanogaster gamma-TuRC is dispensable for targeting gamma-tubulin to the centrosome and microtubule nucleation. J Cell Biol 172, 517–528 (2006).

26. M. Moudjou, N. Bordes, M. Paintrand, M. Bornens, gamma-Tubulin in mammalian cells: the centrosomal and the cytosolic forms. J Cell Sci 109 (Pt 4), 875–887 (1996).

27. S. M. Murphy, L. Urbani, T. Stearns, The mammalian gamma-tubulin complex contains homologues of the yeast spindle pole body components spc97p and spc98p. J Cell Biol 141, 663– 674 (1998).

28. K. Oegema, C. Wiese, O. C. Martin, R. A. Milligan, A. Iwamatsu, T. J. Mitchison, Y. Zheng, Characterization of two related Drosophila gamma-tubulin complexes that differ in their ability to nucleate microtubules. J Cell Biol 144, 721–733 (1999).

29. T. Stearns, M. Kirschner, In vitro reconstitution of centrosome assembly and function: the central role of gamma-tubulin. Cell 76, 623–637 (1994).

30. Y. Zheng, M. L. Wong, B. Alberts, T. Mitchison, Nucleation of microtubule assembly by a gamma-tubulin-containing ring complex. Nature 378, 578–583 (1995).

31. E. Hannak, K. Oegema, M. Kirkham, P. Gonczy, B. Habermann, A. A. Hyman, The kinetically dominant assembly pathway for centrosomal asters in Caenorhabditis elegans is gamma-tubulin dependent. J Cell Biol 157, 591–602 (2002).

32. M. Ohta, Z. Zhao, D. Wu, S. Wang, J. L. Harrison, J. S. Gomez-Cavazos, A. Desai, K. F. Oegema, Polo-like kinase 1 independently controls microtubule-nucleating capacity and size of the centrosome. J Cell Biol 220, (2021).

33. M. D. Sallee, J. C. Zonka, T. D. Skokan, B. C. Raftrey, J. L. Feldman, Tissue-specific degradation of essential centrosome components reveals distinct microtubule populations at microtubule organizing centers. PLoS Biol 16, e2005189 (2018).

34. C. A. Tovey, C. Tsuji, A. Egerton, F. Bernard, A. Guichet, M. de la Roche, P. T. Conduit, Autoinhibition of Cnn binding to gamma-TuRCs prevents ectopic microtubule nucleation and cell division defects. J Cell Biol 220, (2021).

35. K. E. Sawin, P. C. Lourenco, H. A. Snaith, Microtubule nucleation at non-spindle pole body microtubule-organizing centers requires fission yeast centrosomin-related protein mod20p. Curr Biol 14, 763–775 (2004).

36. J. Zhang, T. L. Megraw, Proper recruitment of gamma-tubulin and D-TACC/Msps to embryonic Drosophila centrosomes requires Centrosomin Motif 1. Mol Biol Cell 18, 4037–4049 (2007).

37. Y. K. Choi, P. Liu, S. K. Sze, C. Dai, R. Z. Qi, CDK5RAP2 stimulates microtubule nucleation by the gamma-tubulin ring complex. J Cell Biol 191, 1089–1095 (2010).

38. R. R. Cota, N. Teixido-Travesa, A. Ezquerra, S. Eibes, C. Lacasa, J. Roig, J. Luders, MZT1 regulates microtubule nucleation by linking gammaTuRC assembly to adapter-mediated targeting and activation. J Cell Sci 130, 406–419 (2017).

39. K. W. Fong, Y. K. Choi, J. B. Rattner, R. Z. Qi, CDK5RAP2 is a pericentriolar protein that functions in centrosomal attachment of the gamma-tubulin ring complex. Mol Biol Cell 19, 115– 125 (2008).

40. M. J. Rale, B. Romer, B. P. Mahon, S. M. Travis, S. Petry, The conserved centrosomin motif, gammaTuNA, forms a dimer that directly activates microtubule nucleation by the gamma-tubulin ring complex (gammaTuRC). Elife 11, (2022).

41. Y. Xu, H. Munoz-Hernandez, R. Krutyholowa, F. Marxer, F. Cetin, M. Wieczorek, Partial closure of the gamma-tubulin ring complex by CDK5RAP2 activates microtubule nucleation. Dev Cell 59, 3161–3174 e3115 (2024).

42. M. Serna, F. Zimmermann, C. Vineethakumari, N. Gonzalez-Rodriguez, O. Llorca, J. Luders, CDK5RAP2 activates microtubule nucleator gammaTuRC by facilitating template formation and actin release. Dev Cell 59, 3175–3188 e3178 (2024).

43. M. Wieczorek, T. L. Huang, L. Urnavicius, K. C. Hsia, T. M. Kapoor, MZT Proteins Form Multi-Faceted Structural Modules in the gamma-Tubulin Ring Complex. Cell Rep 31, 107791 (2020).

44. S. Yang, F. K. C. Au, G. Li, J. Lin, X. D. Li, R. Z. Qi, Autoinhibitory mechanism controls binding of centrosomin motif 1 to gamma-tubulin ring complex. J Cell Biol 222, (2023).

45. D. R. Hamill, A. F. Severson, J. C. Carter, B. Bowerman, Centrosome maturation and mitotic spindle assembly in C. elegans require SPD-5, a protein with multiple coiled-coil domains. Dev Cell 3, 673–684 (2002).

46. E. Hannak, M. Kirkham, A. A. Hyman, K. Oegema, Aurora-A kinase is required for centrosome maturation in Caenorhabditis elegans. J Cell Biol 155, 1109–1116 (2001).

47. R. E. Palazzo, J. M. Vogel, B. J. Schnackenberg, D. R. Hull, X. Wu, Centrosome maturation. Curr Top Dev Biol 49, 449–470 (2000).

48. C. A. Kemp, K. R. Kopish, P. Zipperlen, J. Ahringer, K. F. O’Connell, Centrosome maturation and duplication in C. elegans require the coiled-coil protein SPD-2. Dev Cell 6, 511–523 (2004).

49. L. Pelletier, N. Ozlu, E. Hannak, C. Cowan, B. Habermann, M. Ruer, T. Muller-Reichert, A. A. Hyman, The Caenorhabditis elegans centrosomal protein SPD-2 is required for both pericentriolar material recruitment and centriole duplication. Curr Biol 14, 863–873 (2004).

50. M. Boxem, Z. Maliga, N. Klitgord, N. Li, I. Lemmens, M. Mana, L. de Lichtervelde, J. D. Mul, D. van de Peut, M. Devos, N. Simonis, M. A. Yildirim, M. Cokol, H. L. Kao, A. S. de Smet, H. Wang, A. L. Schlaitz, T. Hao, S. Milstein, C. Fan, M. Tipsword, K. Drew, M. Galli, K. Rhrissorrakrai, D. Drechsel, D. Koller, F. P. Roth, L. M. Iakoucheva, A. K. Dunker, R. Bonneau, K. C. Gunsalus, D. E. Hill, F. Piano, J. Tavernier, S. van den Heuvel, A. A. Hyman, M. Vidal, A protein domain-based interactome network for C. elegans early embryogenesis. Cell 134, 534–545 (2008).

51. M. Decker, S. Jaensch, A. Pozniakovsky, A. Zinke, K. F. O’Connell, W. Zachariae, E. Myers, A. A. Hyman, Limiting amounts of centrosome material set centrosome size in C. elegans embryos. Curr Biol 21, 1259–1267 (2011).

52. A. Thawani, S. Petry, Molecular insight into how gamma-TuRC makes microtubules. J Cell Sci 134, (2021).

53. D. K. Dhani, B. T. Goult, G. M. George, D. T. Rogerson, D. A. Bitton, C. J. Miller, J. W. Schwabe, K. Tanaka, Mzt1/Tam4, a fission yeast MOZART1 homologue, is an essential component of the gamma-tubulin complex and directly interacts with GCP3(Alp6). Mol Biol Cell 24, 3337–3349 (2013).

54. T. L. Huang, H. J. Wang, Y. C. Chang, S. W. Wang, K. C. Hsia, Promiscuous Binding of Microprotein Mozart1 to gamma-Tubulin Complex Mediates Specific Subcellular Targeting to Control Microtubule Array Formation. Cell Rep 31, 107836 (2020).

55. N. Janski, K. Masoud, M. Batzenschlager, E. Herzog, J. L. Evrard, G. Houlne, M. Bourge, M. E. Chaboute, A. C. Schmit, The GCP3-interacting proteins GIP1 and GIP2 are required for gamma-tubulin complex protein localization, spindle integrity, and chromosomal stability. Plant Cell 24, 1171–1187 (2012).

56. T. C. Lin, A. Neuner, D. Flemming, P. Liu, T. Chinen, U. Jakle, R. Arkowitz, E. Schiebel, MOZART1 and gamma-tubulin complex receptors are both required to turn gamma-TuSC into an active microtubule nucleation template. J Cell Biol 215, 823–840 (2016).

57. H. Munoz-Hernandez, Y. Xu, A. Pellicer Camardiel, D. Zhang, A. Xue, A. Aher, E. Walker, F. Marxer, T. M. Kapoor, M. Wieczorek, Structure of the microtubule-anchoring factor NEDD1 bound to the gamma-tubulin ring complex. J Cell Biol 224, (2025).

58. M. Nakamura, N. Yagi, T. Kato, S. Fujita, N. Kawashima, D. W. Ehrhardt, T. Hashimoto, Arabidopsis GCP3-interacting protein 1/MOZART 1 is an integral component of the gamma-tubulin-containing microtubule nucleating complex. Plant J 71, 216–225 (2012).

59. M. Wurtz, E. Zupa, E. S. Atorino, A. Neuner, A. Bohler, A. S. Rahadian, B. J. A. Vermeulen, G. Tonon, S. Eustermann, E. Schiebel, S. Pfeffer, Modular assembly of the principal microtubule nucleator gamma-TuRC. Nat Commun 13, 473 (2022).

60. T. C. Lin, A. Neuner, Y. T. Schlosser, A. N. Scharf, L. Weber, E. Schiebel, Cell-cycle dependent phosphorylation of yeast pericentrin regulates gamma-TuSC-mediated microtubule nucleation. Elife 3, e02208 (2014).

61. J. M. Kollman, J. K. Polka, A. Zelter, T. N. Davis, D. A. Agard, Microtubule nucleating gamma-TuSC assembles structures with 13-fold microtubule-like symmetry. Nature 466, 879–882 (2010).

62. A. F. Brilot, A. S. Lyon, A. Zelter, S. Viswanath, A. Maxwell, M. J. MacCoss, E. G. Muller, A. Sali, T. N. Davis, D. A. Agard, CM1-driven assembly and activation of yeast gamma-tubulin small complex underlies microtubule nucleation. Elife 10, (2021).

63. C. Frokjaer-Jensen, M. W. Davis, C. E. Hopkins, B. J. Newman, J. M. Thummel, S. P. Olesen, M. Grunnet, E. M. Jorgensen, Single-copy insertion of transgenes in Caenorhabditis elegans. Nat Genet 40, 1375–1383 (2008).

64. A. Dammermann, T. Muller-Reichert, L. Pelletier, B. Habermann, A. Desai, K. Oegema, Centriole assembly requires both centriolar and pericentriolar material proteins. Dev Cell 7, 815–829 (2004).

